# Sustained High- and Low-Frequency Neural Responses to Visual Stimuli in Scalp EEG

**DOI:** 10.1101/290593

**Authors:** Edden M. Gerber, Leon Y. Deouell

**Affiliations:** Hebrew University of Jerusalem

## Abstract

What are the neurophysiological correlates of sustained visual processing in the scalp EEG signal? In a previous study using intracranial recordings in humans, we found that presentation of visual stimuli for prolonged durations (up to 1.5 seconds) was associated with two kinds of sustained neural activity patterns: a high-frequency broadband (>30 Hz) response that tracked the duration of the stimulus with high precision in early visual cortex (EVC), and with lesser temporal precision in downstream category-selective areas; and a sustained low-frequency potential shift appearing in a small subset of EVC sites. Using a similar approach of presenting images for variable durations to identify sustained activity, we provide the first comprehensive characterization of the manifestation of sustained visual responses as recorded with EEG. In a series of four experiments, we found that both high- and low-frequency sustained responses can be detected on the scalp. The high frequency activity could be detected with high signal to noise ratio only in a subset of individual subjects, in whom it was unequivocal and highly localized. The low frequency sustained response was sensitive to the size and position of the stimulus in the visual field. Both response types showed strong lateralization for stimuli on the left vs. right visual field, suggesting a retinotopic visual cortical source. However, different scalp topographies and different modulation by stimulus properties suggest that the two types of sustained responses are likely driven by distinct sources, and reflect different aspects of sustained processing in the visual cortex.

## Introduction

The neural mechanisms underlying vision are commonly investigated by measuring neural responses to artificial visual stimuli. These stimuli are typically presented for short durations, for the sake of efficient data collection. As a result, the observed neural responses are driven mostly by events of change in the visual field (e.g. appearance or disappearance of the stimulus), whereas the ongoing component of vision – that is, the neural activity associated with continuously perceiving an image in the visual field, rather than with moments of sudden change – is left unaccounted for. This reduction of the space of visual experience to briefly-presented stimuli may stem from the notion, characteristic of electrical engineering, that the impulse response of a system (the response to a very short stimulus, avoiding overlap of responses) is a good way for characterizing it. In a previous study (Gerber et al., 2017), we employed visual stimuli of variable duration in an intracranial study in humans and found that ongoing activity associated with stimulus presence and persisting for the entire duration of stimulus presentation (“duration-tracking” responses) are found in the broadband high-frequency (>30Hz) signal across the visual cortex. This sustained response showed a posterior to anterior gradient: it was robust and reliable in the early visual cortex (EVC), whereas sustained activity in high-level visual areas was present on average but was weaker and much less reliable than in EVC on a single-trial level. In addition, we found that a subset of electrodes in the early visual cortex exhibited a duration-tracking sustained positive or negative shift of the local field potential (LFP), lasting until ∼200 ms after stimulus offset. This low frequency shift did not necessarily co-occur in the same electrodes as the high-frequency sustained response.

In the present exploratory study, we investigated duration-tracking neural responses to static, variable duration stimuli in scalp EEG, focusing on high frequency (i.e. “gamma” range) power and (lower frequency) event related potentials. High-frequency activity on the scalp has been widely studied as a marker of visual processing in various contexts (Adjamian et al., 2004; Busch et al., 2006; Hall et al., 2005; Hoogenboom et al., 2006; Perry et al., 2013; Swettenham et al., 2009; Tallon-Baudry et al., 2005; Tallon-Baudry and Bertrand, 1999), but no studies to our knowledge have attempted to characterize the sustained high frequency response component as distinct from onset- or motion-driven activity. Additionally, the presence of ocular and myogenic noise sources presented major challenges for the detection and interpretation of this type of activity (Muthukumaraswamy, 2013; Rice et al., 2013; Whitham et al., 2008; Yuval-Greenberg et al., 2008). While the onset of a visual stimulus may be associated with a number of non-neural sources of noise that may contribute to the recorded response (e.g. changes in the saccade rate or a motor response), these noise sources are less likely to be contingent on the time course of the presence of the stimulus, and so duration tracking in the high-frequency response would serve as strong evidence that this signal is indeed neural in origin.

Sustained modulations of the evoked potential in response to prolonged stimuli have been demonstrated in the auditory domain, where the evolution of stimuli in time is a cardinal aspect of the auditory information. In that modality, it was shown that a continuous acoustic signal produces a sustained baseline shift in the scalp EEG signal (Hillyard and Picton, 1978; Pantev et al., 1994; Picton et al., 1978). In the visual domain, a sustained baseline shift in the EEG and MEG signal was demonstrated in response to a simple variable duration stimulus (a green LED light), though only in the context of an explicit duration estimation task (Maquet et al., 1996; N’Diaye et al., 2004; Pouthas et al., 2000). To our knowledge, no attempt has been made to characterize the sustained component of the visual evoked response as a neural correlate of ongoing visual perception distinct from activity driven by changes in the visual field and from the requirement to explicitly estimate the stimulus duration.

Using variable-duration stimuli is necessary for identifying the sustained response component that is contingent on stimulus presence and separating it from the onset response. The onset-driven response elicited by a fixed-duration stimulus may persist beyond the actual onset, and if the duration of this onset response happens to equal the duration of the stimulus, it would be erroneously dimmed to be a duration-tracking response. In contrast, any difference in the duration of the responses associated with stimuli of at least two different durations can be reliably attributed to the continued presence of the stimulus. Using very long stimulus durations (e.g. several seconds) and showing that the neural response returns to baseline shortly after the stimulus offset can also provide convincing evidence of duration tracking activity (see for example Gilaie-Dotan et al., 2008 showing sustained fMRI BOLD responses), however this is not a time-efficient method of identifying such responses. Additionally, a distinction is necessary between variable-duration static image stimuli and moving stimuli (e.g. movie clips, rotating shapes etc.): while the latter are in some cases optimal in driving neural activity (Hoogenboom et al., 2006; Muthukumaraswamy and Singh, 2013), the fact that they are dynamic means that they cannot be used to dissociate activity associated with ongoing perception to that associated with the response to changes in the visual field.

In a series of four experiments, we varied durations and stimuli, aiming to characterize the sustained high frequency and low frequency responses of the EEG, along two main dimensions: retinotopic position and semantic content. We report that highly localized broadband high frequency responses can be recorded on the human scalp of single subjects, albeit only in a subset of the subjects. The ERP sustained response is more widely distributed and more ubiquitous across subjects.

## Materials and Methods

### Subjects

Sixty-two healthy subjects in total participated in the four experiments. Thirteen subjects participated in Experiment 1 (7 female, ages 18-30), 12 in Experiment 2 (8 female, ages 18-51), 14 in Experiment 3 (8 female, ages 19-34) and 30 in Experiment 4 (19 female, ages 18-27). Two of the subjects participated in three experiments, and an additional three subjects participated in two. All subjects had normal or corrected to normal vision. Subjects provided written informed consent and received either payment (∼$10 per hour) or course credit for participation in the experiment. The study was approved by the Ethics Committee of the Hebrew University of Jerusalem.

### Experimental procedures

In all experiments, subjects were seated in a dimly lit, sound attenuated double wall chamber (Eckel C-26; Eckel, UK), 100 cm in front of a cathode ray tube (CRT) monitor (Viewsonic, 100Hz refresh rate) with their heads supported by a chin and forehead rest.

#### Experiment 1

The goal of experiment 1 was to identify low frequency baseline shift and high frequency duration-tracking visual activity based on EEG responses to visual stimuli of variable duration, using the same material used in the study of Gerber et al. (2017) in which duration-tracking responses were recorded intracranially. The experiment consisted of 1300 trials, divided into 20 recording blocks of 2.5 minutes each. In each trial, subjects were presented with a central rectangular grayscale image of a face, object, body part, animal, or clothing item, extending 3.1 visual degrees to each side of the center of the screen (Figure 1A). The stimuli were presented for a duration of either 300, 900 or 1500 milliseconds, with an inter-stimulus-interval (ISI) of between 600 and 2400 ms (in 450 ms increments) including a central fixation cross. The total duration of each trial was therefore between 900 and 3900 ms. Because a minimal trial duration of 1800 ms (accommodating a 1500 ms stimulus plus an additional 300 ms post-offset period) was required for comparing trials of different durations, 300 trials with a total duration (stimulus + ISI) of less than 1800 ms were excluded from analysis. Subjects were asked to maintain fixation on the fixation cross or on the center of the presented image and to respond with a button press to the images of clothing items (10% of the trials). These target trials were also excluded from analysis, leaving 300 trials for each stimulus duration.

**Figure 1:**
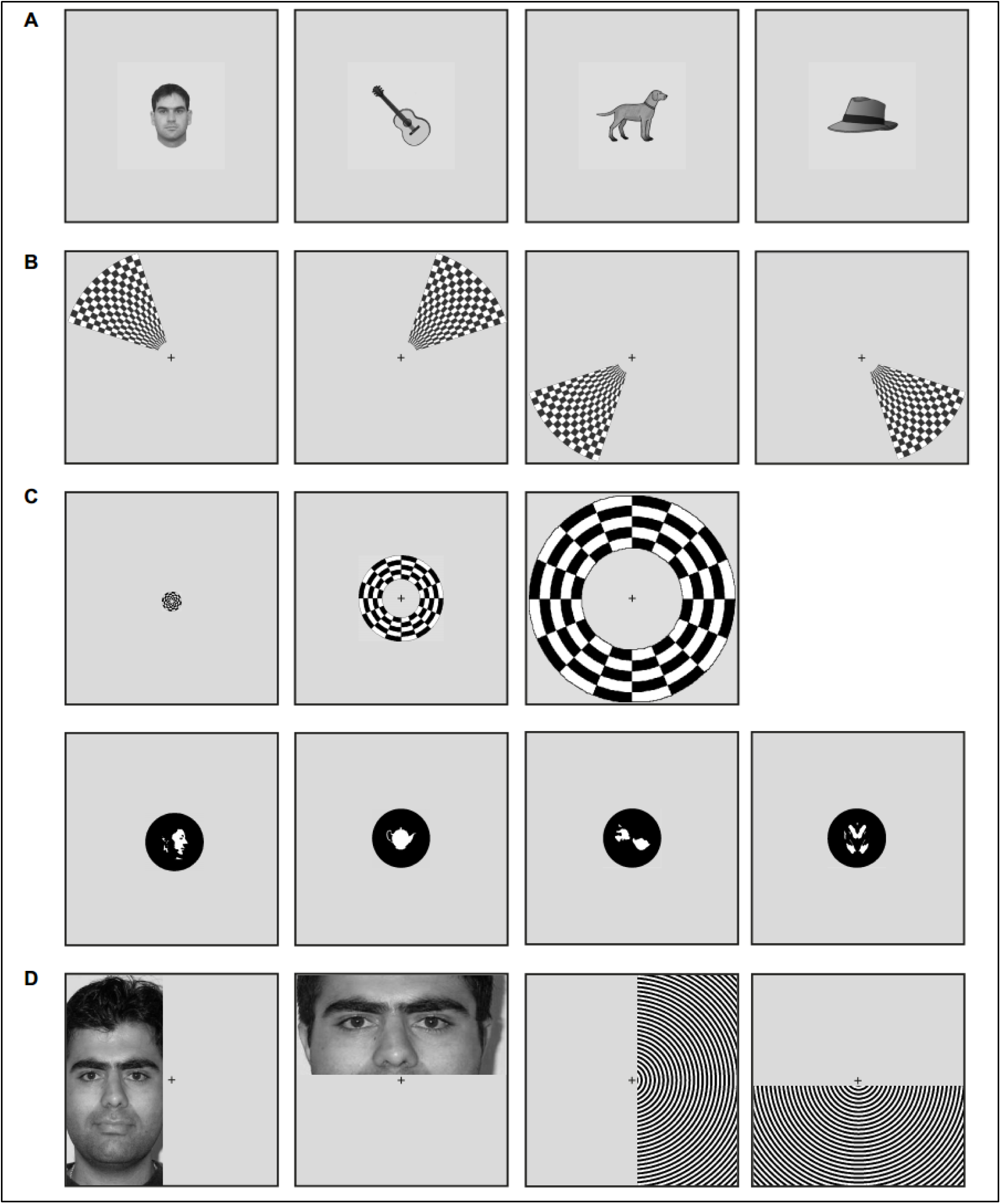
Stimuli in experiments 1-4. Each panel shows several example stimuli from the four experiments EEG included in this study. Each stimulus is presented to scale within a square that corresponds to the monitor display (the actual display had additional margins on the left and right due to the rectangular shape of the monitor). **(A)** Experiment 1. Left to right: three example stimuli (face, object, animal) and an example target stimulus (clothing items). **(B)** Experiment 2. Stimuli were checkerboard wedges appearing in one of four corners of the display. Target stimuli were a subset of words within a stream of words appearing over the fixation cross. **(C)** Experiment 3. Top row, left to right: checkerboard ring stimuli with small, medium, and large eccentricity. Bottom row, left to right: example stimuli from the Mooney face, object, and scrambled conditions, and a target stimulus consisting of a symmetric scrambled image. **(D)** Experiment 4. Stimuli were either faces or circular grating stimuli, appearing in either the left, right, top or bottom half of the display.

#### Experiment 2

Experiment 2 was designed to test the effect of the stimulus’ vertical and horizontal position on the duration-tracking high frequency and baseline shift responses. The experiment consisted of 1333 trials in 18 recording blocks of about 3.3 minutes each. Stimuli were black and white checkerboard wedges appearing in the top right, top left, bottom right or bottom left of the screen (Figure 1B). The wedges extended from 1 to 6.4 visual degrees of eccentricity, with a polar angle of 54 degrees, and were divided into 16 checkerboard sections on the eccentricity axis and 15 sections on the polar axis. Stimuli were presented for 300, 900 or 1500 ms as in experiment 1. The inter-stimulus interval was 1500 to 2100 ms in 150 ms increments, such that the minimal trial duration was 1800 ms. To engage the subjects on the task and help the subject maintain central fixation, they were asked to fixate on a central fixation cross while words appeared one at a time over the cross, and to respond with a button press when the word was a name of an animal. Words appeared every 7.8 sec on average for 300 ms each. 133 trials in which a word appeared within the 1800 ms post-stimulus-onset window were excluded, leaving 1200 trials for analysis. Stimulus position and duration were randomized across trials, but the 1200 analyzed trials always consisted of 100 trials from each condition (duration and position combination).

#### Experiment 3

Experiment 3 was designed to test two separate issues: the effect of stimulus eccentricity and the effect of semantic content on the duration-tracking baseline shift. The experiment consisted of 1333 trials in 20 recording blocks of about 2.5 minutes each. The stimuli were either a black and white checkerboard ring of one of three possible sizes, or a central black-and-white circular image (with black background). The image contained either a “Mooney face” image (pictures of human faces where all shades are replaced with either black or white, thus leaving a pattern of patches evoking the percept of a face), a “scrambled” Mooney face where the white patches were rotated and shifted so that no recognizable shape appeared, or a white silhouette of an object (Figure 1C). The stimuli were presented for either 500 or 1000 ms, with an ISI of 800 to 1400 ms in 150 ms increments, for a total trial duration of 1300 to 2400 ms. Subjects were required to fixate on a central fixation cross and to respond with a button press to target stimuli, which were scrambled Mooney faces arranged in a symmetric pattern (10% of trials). These target trials were excluded from analysis, leaving 1200 trials divided into 100 trials from each condition (duration and stimulus type). The checkerboard rings extended from either 0.15 to 0.75 (small), 1.5 to 3.3 (medium), or 4 to 8.1 (large) visual degrees of eccentricity, and were divided into 5 checkerboard sections on the radial axis and 16 on the polar axis. These eccentricity ranges were selected to maintain the estimated area of cortical activation as equal as possible between the three conditions, in accordance with the increasing cortical magnification along the visual eccentricity axis. Cortical magnification is commonly modeled by the formula:

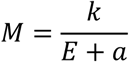

Where *M* is the magnification factor in mm/degree, *E* is the eccentricity in degrees, and *k* and *a* are fitted parameters (Schira et al., 2007). To estimate the relative cortical magnification for the area activated by each pair of checkerboard rings, published data from Schira and colleagues, covering the 0.75-12 degree range (Schira et al., 2007), and the 0-4.8 degree range (Schira et al., 2009), was pooled together and new parameters for the above formula were fitted to it (Figure S1). Based on these parameters and the formula for cortical magnification, the cortical area activated by two rings of different eccentricities could be equalized with the equation

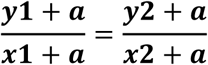

Where *x1* and *y1* are the inner and outer radius of the first ring, *x2*, *y2* are the inner and outer radius of the second ring, and *a* is the fitted parameter. Throughout the experiment, subjects were required to fixate on a fixation cross appearing in the center of the screen.

#### Experiment 4

Experiment 4 was designed to test the modulation of visual duration-tracking responses by the stimulus horizontal and vertical position, as well as to test whether simple (grating) and complex (face) stimuli will produce a different topography of high frequency activity on the scalp, due to activity in high-level visual areas. The experiment consisted of 1456 trials divided into 20 blocks of about 2.3 minutes each. The stimuli were rectangular images appearing in either the top, bottom, left or right half of the screen, starting 0.2 visual degrees away from the center and extending 6.4 degrees toward the edge of the screen (Figure 1D). The images were either the respective top, bottom, left or right half of a black and white image of a circular grating, or a grayscale face image (showing a full face for the vertically-oriented left/right stimuli, and a partial image centered around the eyes and nose for horizontally-oriented top/bottom stimuli). For the grating stimuli, full-contrast black and white annular square grating stimuli with a 3 cycles-per-visual-degree spatial frequency were selected as an optimal stimulus for generating high-frequency neural responses (Adjamian et al., 2004; Perry et al., 2015). Stimulus durations were 300 or 900 ms with an ISI of 1100 to 1400 ms in 100 ms increments, for a total trial duration of 1400 to 2100 ms. Subjects were required to maintain fixation on a central fixation cross throughout each experiment block. Trials in which subjects’ gaze deviated more than 0.2 degrees from the fixation on average throughout the trial were excluded from analysis. The subjects’ task was to respond with a button press whenever a stimulus did not disappear at once but with a 200-ms gradual fade, requiring subjects to attend to the stimuli throughout their presentation duration. 11% of the trials were target trials that were not analyzed, leaving 1296 trial for analysis, divided into 81 trials from each experimental condition (combination of stimulus type, position and duration).

### EEG recording

EEG was recorded with an Active 2 acquisition system (Biosemi, the Netherlands) using sintered Ag/AgCl electrodes, at a sampling rate of 1024 Hz with online low-pass filter at 204.8 Hz to prevent aliasing of high frequencies. A 64-electrode cap organized according to the extended 10-20 system was used in all experiments (http://www.biosemi.com/pics/cap_64_layout_medium.jpg). Eight additional electrodes were placed: two on the mastoid processes, two horizontal EOG electrodes lateral to the left and right eyes’ lateral canthi, three vertical EOG electrodes below (infraorbital) and above (supraorbital) the right eye, and below the left eye, and an electrodes on the tip of the nose. All electrodes were referenced during recording to a common-mode signal (CMS) electrode between POz and PO3, and were subsequently digitally re-referenced to the average of the scalp electrodes (see Data Processing below).

### Eye-movement recording

Binocular eye movements were monitored in all four experiments using an Eyelink 1000/2K infrared video-oculographic desktop system (SR Reasearch Ltd., Ontario, Canada). Left and right eye position data were recorded continuously at a rate of 1000 Hz and saved for off-line analysis. Triggers sent via a parallel port from the stimulation computer were split and recorded simultaneously by the EEG recording system and by the eye tracker and were used to synchronize the EEG and eye tracking recording. To correct for apparent shifts of gaze position caused by changes in pupil size during the experiment (on the scale of ∼1 visual degree (Drewes et al., 2012)) we applied a regression analysis to individual subjects’ vertical and horizontal gaze position with pupil size serving as the regressor, and subtracted the time course predicted by this factor.

### Data processing

Data processing was done with custom Matlab code (Mathworks, Natick, MA), with some functionality adopted from the Fieldtrip toolbox (Oostenveld et al., 2011) and the EEGLab toolbox (Delorme and Makeig, 2004). EEG data from all experiments was down-sampled to 512 Hz and re-referenced to the average of the 64 scalp electrodes (excluding occasional electrodes determined to include excessive noise across the entire experiment duration based on visual inspection). Each continuous recording block was individually de-trended by subtracting the linear vector connecting the means of the initial and final 10 samples of the block to avoid edge artifacts during filtering, and the de-trended blocks were concatenated back together. The data was high-pass filtered with a cutoff of 0.1 Hz with a 3^rd^ degree Butterworth filter, and 50Hz line noise was removed with a custom notch filter designed to suppress continuous 50 Hz oscillations, typical of power lines noise, while having less effect on more transient components (Keren et al., 2010). To attenuate noise driven by eye movements and muscle activity, independent component analysis (ICA) was applied as described in (Keren et al 2010). ICA training data was generated by concatenating all post-stimulus segments as well as 60 ms segments centered on every saccade-onset event detected algorithmically based on eye-tracker data (Engbert, 2006; Keren et al., 2010). ICA components were manually identified as reflecting ocular or myogenic noise based on their temporal profile, scalp topography, power spectrum, and low-frequency power modulation around stimulus onset. To remove noise components, these components were multiplied by zero and the data was then re-mixed into channel data. Next, epochs including extreme-value artifacts were identified. For the ERP analysis, artifacts were defined as peri-stimulus segments exceeding a threshold value of 100 µV. The data was segmented around stimulus onsets with a 300 ms pre-stimulus baseline window. The post-stimulus window length varied for each experiment’s, ending 300 or 500 ms after the longest duration. Each trial was baseline-corrected by subtracting the mean of the 300 ms pre-stimulus window from all data points, and trials were averaged (excluding trials overlapping with artifacts) to produce the ERP for each condition.

We computed the high frequency power signal as the 50-100Hz band-limited power modulation by filtering the EEG in this band using a 3^rd^ degree band-pass Butterworth zero phase shift filter, and taking the square of the Hilbert transform of the filtered data. Artifacts in this measure were defined as above a threshold of 50 µV^2^. Finally, trials were rejected if a blink occurred within the initial 300 ms of stimulus presentation. The band-limited power signal was segmented around stimulus onsets similarly to the ERP data, baseline-corrected by subtracting the mean of the 300 ms pre-stimulus window and averaged after discarding trials overlapping with high-frequency artifacts. Morlet wavelet analysis was also performed in selected cases for visualization purposes only (Figures 4, 6, 8, S3), with a frequency range of 2-160Hz (2Hz increments) and a wavelet constant of 12. Wavelet-transformed data was segmented and baseline-corrected by subtracting the mean pre-stimulus amplitude separately for each frequency.

### Duration-tracking score

The goal of the duration-tracking analysis was to identify electrodes where longer stimulus durations elicit correspondingly longer responses. Consider two stimuli, one of duration X ms, and the other of duration X+Y ms. Under H1 of a sustained response matching the duration of the stimuli, we expect the response to the two stimuli to be similar in the first X ms, but to differ in the next Y ms (Figure 2). Therefore, to test for duration-tracking, responses to two different stimulus durations were compared using a timepoint-by-timepoint t-test, resulting with a vector of t-values. This comparison was done either across single trials for a single-subject analysis or across subject averages for the group-average analysis. A threshold was applied to the t-value vector corresponding to a p-value of 0.01 (uncorrected), and supra-threshold t-values (either above the positive threshold or below the negative threshold) were summed across a temporal window in which, under H1, duration-tracking responses should differ. This window started at the end of the shorter stimulus duration, and ended 300 ms following the end of the longer duration (e.g., 300-1200 ms when comparing 300 vs 900 ms stimulus durations). This value was then divided by the actual temporal difference between the two stimulus durations (in samples) to yield a sample-wise normalized tracking score (Figure 2). In cases where the experiment included three stimulus durations, this value was computed for each pair of consecutive durations (e.g. 300 vs 900 ms, and 900 vs 1500 ms) and the average of the two tracking scores was used. The resulting duration-tracking score thus reflects the mean significant t-value of the difference between the two responses within the critical sustained response epoch. The procedure is derived from the cluster-based permutation test (Maris and Oostenveld, 2007), except that there is no requirement of above-threshold time points to be contiguous (we relaxed this requirement to better capture a sustained difference across the entire analyzed window, rather than a temporally-localized cluster of difference). Figure 2 shows that the contribution of offset responses which necessarily occur at different times for stimuli of different durations, is typically canceled out. This is due to the fact these responses appear in opposite polarities in the t-value vector comparing the two durations (Figure 2B, bottom), and thus negative and positive t-values within the analyzed time window offset one another (to the degree that the offset responses are of similar magnitude). The supra-threshold t-values were used, as opposed to averaging all t-values within the window, to avoid interpreting very small but consistent differences between responses as large effects. To derive a statistical significance estimate for this measure, the observed duration-tracking score was compared against a null-hypothesis distribution generated by a 10,000-iteration permutation test where the stimulus duration labels were randomly shuffled. Duration-tracking was tested separately in each electrode and stimulus condition. A Bonferroni correction for multiple comparisons was applied separately for each of these tests, dividing the 0.05 significance threshold by the number of electrodes (72 in the first experiment, and 19 in subsequent experiments following the definition of a posterior region-of-interest).

**Figure 2:**
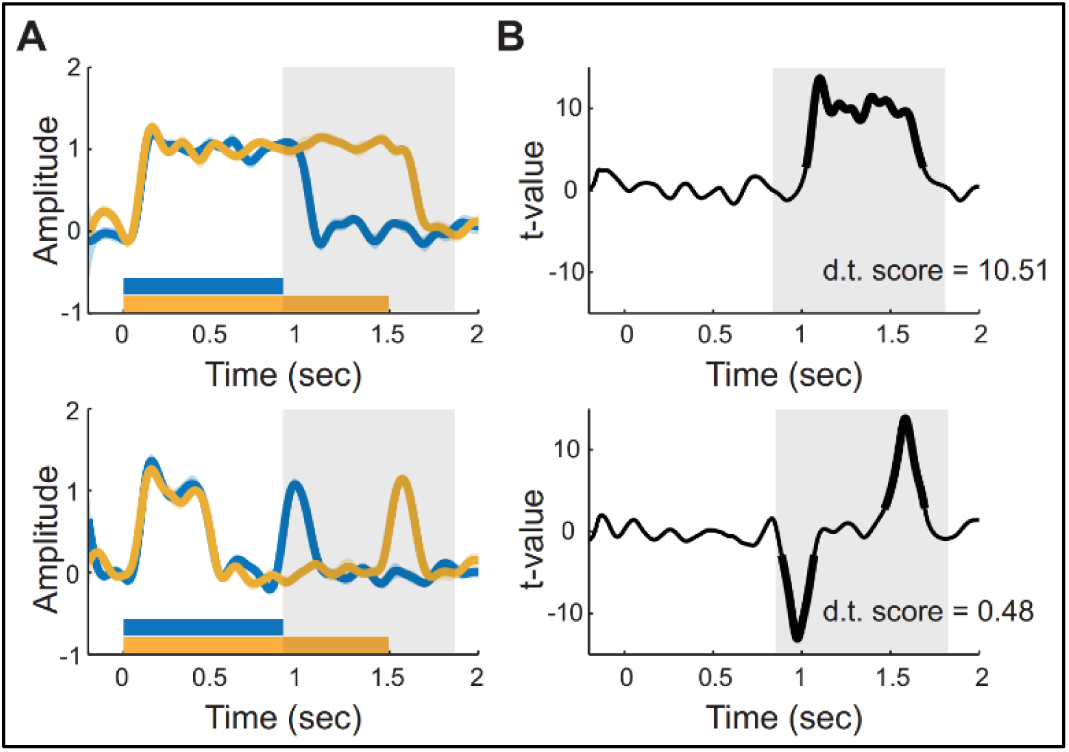
Computing the duration-tracking score. The duration-tracking score reflects the degree to which the neural response is sustained for the entire stimulus duration. **(A)** Schematic examples of duration-tracking response (top) and non-duration-tracking response (bottom) to a 0.9 and a 1.5 sec stimulus (bars at bottom of each plot correspond to stimulus duration). Note that the non-duration-tracking example contains offset components which are dependent on stimulus duration, but should not be considered duration-tracking as they are not sustained. **(B)** Duration-tracking score is computed by deriving a point-by-point t-value of the difference between the responses to the two durations, and summing all supra-threshold values within the time window where the difference is expected (0.9 to 1.8 sec in this example, shown by gray rectangle; thick traces correspond to supra-threshold t-values). This value is then normalized by the time difference.

## Results

### Experiment 1

Thirteen subjects were shown images of faces and objects for either 300, 900 or 1500 ms, and pressed a key in response to a target image category (clothes). All subjects performed the task successfully (hit rate: mean 98.5%, std. deviation 2.3%; false alarms: mean 1%, std. deviation 0.3%).

#### Duration-tracking in the event related potential

The event-related potential in posterior electrodes exhibited a robust duration-tracking response, manifesting as a sustained positive plateau-like shift that returned to baseline approximately 300 ms after the stimulus offset (Figure 3A,B). Applying the duration-tracking analysis (see Materials and Methods) revealed a significant effect with a broad posterior topography, appearing in a symmetric 12-electrode posterior cluster in the group average, with a maximum at electrode Iz (permutation p<0.05, Bonferroni-corrected for 72 electrodes). Applying the same metric at the individual subject level (with variance across trials) revealed a significant effect in at least one posterior electrode in eleven of the thirteen subjects (figure 3C,D).

**Figure 3:**
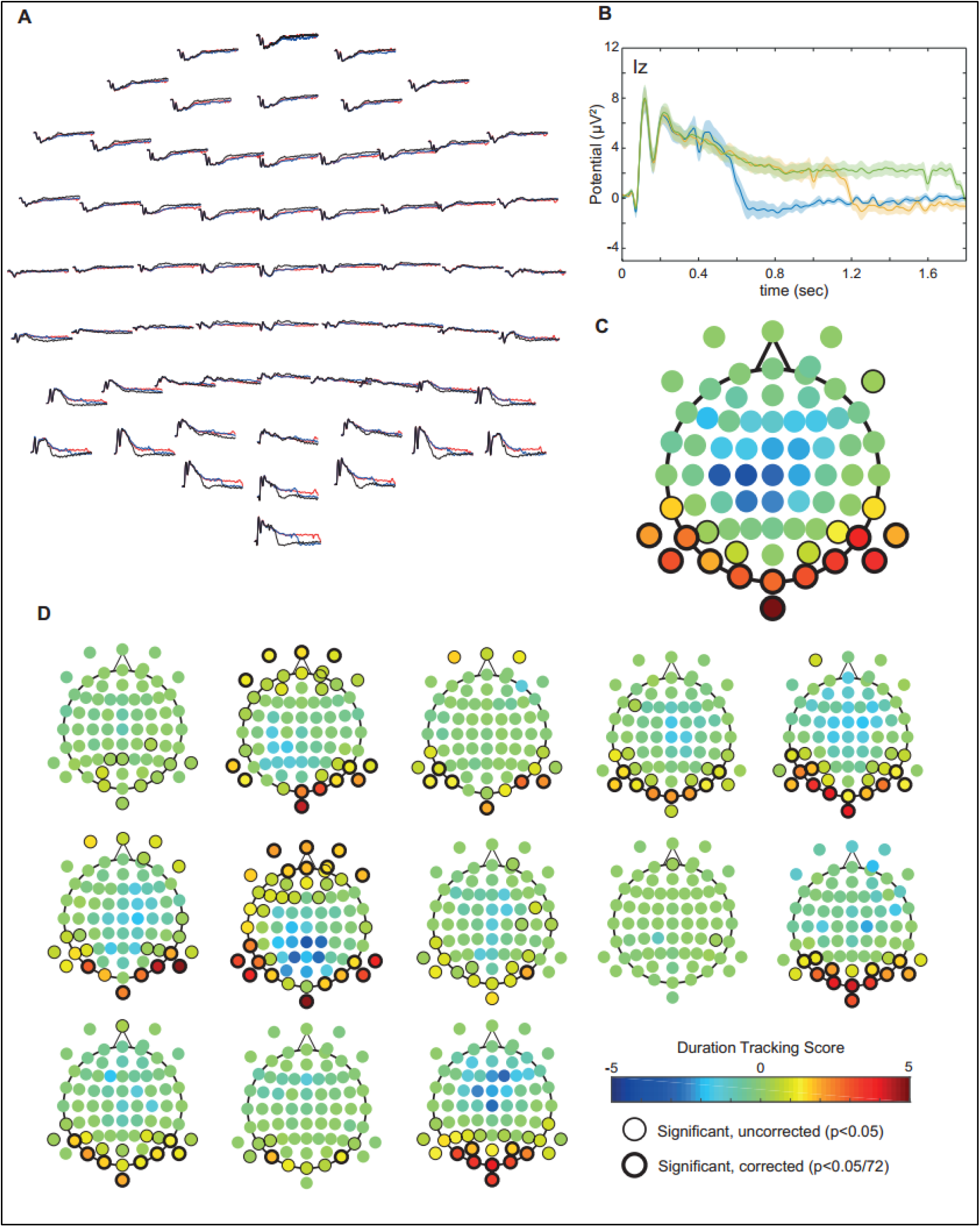
Experiment 1 – duration-tracking visual baseline shift. **(A)** Event-related responses to visual stimuli displayed for 300, 900 and 1500 ms, averaged across subjects and shown for every scalp electrode. **(B)** Same as (A), showing only Iz. The shaded area around the lines represent the standard error across subjects **(C)** Scalp topography of the duration-tracking magnitude in the subject average. Legend and color bar are as in panel D. **(D)** Scalp topography of the duration-tracking score and permutation-based significance in thirteen individual subjects.

#### Duration-tracking in the high frequency power

The same analysis was applied to the high frequency band-limited (50-100Hz) power signal. Significant duration-tracking high-frequency responses were found in three subjects (Figure 4A). Wavelet analysis of the data between 2-160 Hz revealed in these subjects a sustained wideband increase in spectral power compared to baseline levels between about 50Hz and at least 120Hz. This response has a consistent, highly localized right posterior topography in all 3 subjects (with a maximum at either one of the adjacent electrodes O2 and PO8), with unequivocal correspondence to stimulus duration. This duration-tracking was observed for the 50-100Hz band-limited power signal, as well as in individual narrow frequency bands up to about 120Hz (indicating that the sustained response is indeed broadband in nature). Two additional subjects showed a highly similar response in the same region that did not reach the significance threshold after Bonferroni correction. The duration-tracking effect was too sparse across subjects to reach significance in the group average; however, the magnitude of the duration-tracking effect in the group average exhibited the same right-posterior topography (Figure 4B).

**Figure 4:**
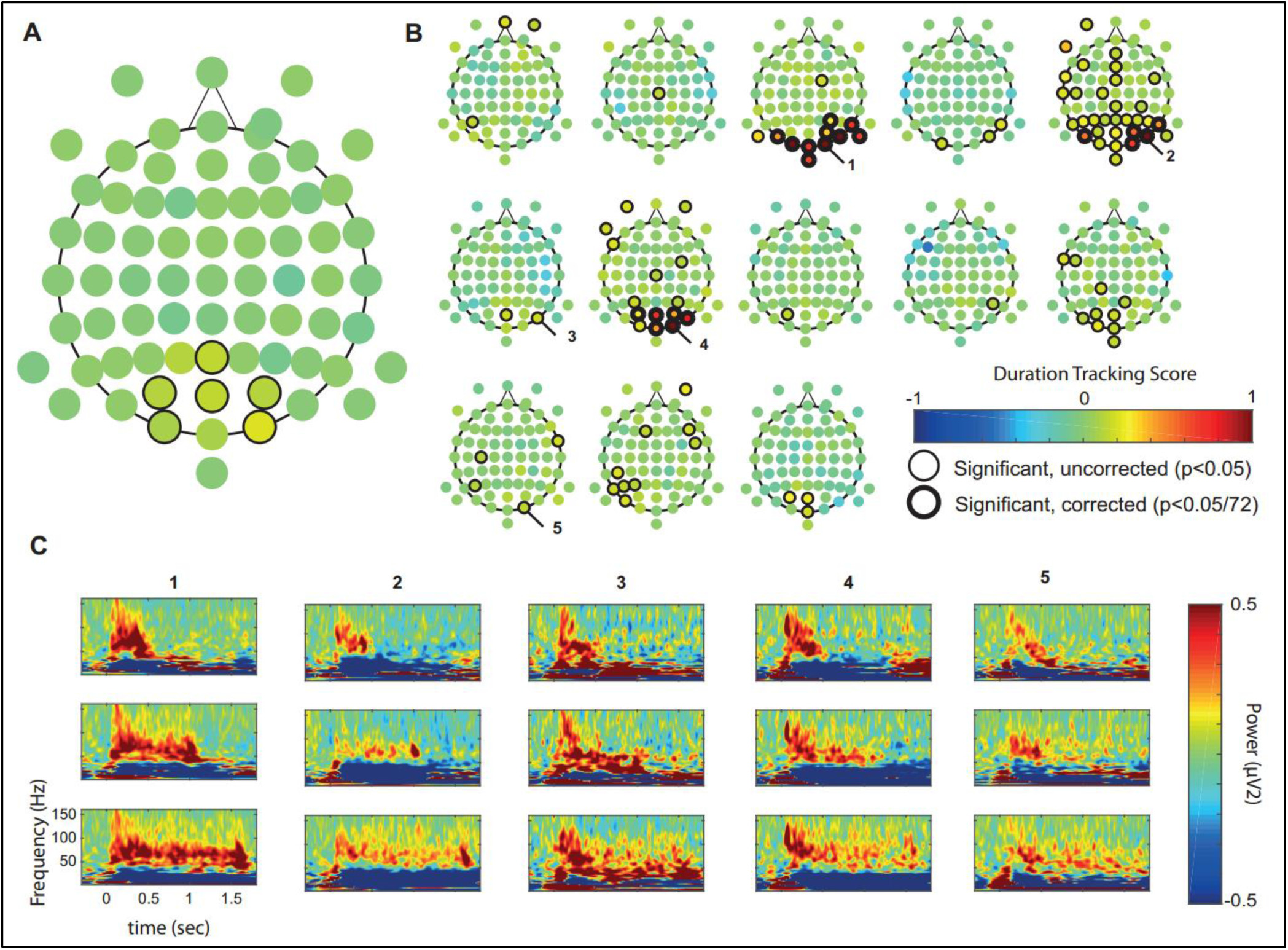
Experiment 1 – duration-tracking visual high-frequency responses. **(A)** Scalp topography of the duration-tracking magnitude of the high frequency band-limited power signal in individual subjects. **(B)** Scalp topography of the effect in the subject average. While not significant after Bonferroni correction, effect size is largest in electrode O2. **(C)** Mean time-frequency plots of high-frequency responses in individual subjects, in the electrodes marked with corresponding numbers in panel A. Each electrodes’ plots from top to bottom correspond to 300 ms, 900 ms, and 1500 ms stimulus.

#### Eye movements do not drive the duration-tracking baseline shift or high frequency response

A sustained activity profile in response to the prolonged presence of a visual stimulus could potentially be the result of averaging across trials containing multiple transient responses at random time points, specifically those driven by eye movements. In addition to the myogenic saccadic spike potential (Yuval-Greenberg et al., 2008), saccadic movements occurring at random latencies throughout stimulus presentation could elicit transient potentials (known as lambda waves) and/or bursts of high-frequency activity in the visual cortex, driven by the change in the input to the visual system. These could then appear as a sustained response after averaging across trials. To test this possibility, we examined individual subjects’ trials showing different rates of saccades (and microsaccades) as detected in the high resolution eye-tracker data. Specifically, we sorted the 0.9 sec stimulus duration trials by the number of saccades occurring within a 1.2-second window following stimulus onset (i.e. from its onset to 300 ms after the offset). In each subject, we grouped together all trials that included no saccades at all in this window (no saccades condition), and the same number of trials which had the highest number of saccades (high saccade rate condition). To test the effect of saccades on the sustained low frequency event related potential, we tested the difference between the responses in the no saccade and high saccade rate responses across subjects in electrode Iz, where the duration-tracking baseline shift effect was maximal. The responses were very similar and a cluster-based permutation test (Maris and Oostenveld, 2007; 10,000 permutations) revealed no significant differences (Figure 5A). To test the effect of saccades on the high-frequency sustained response, a similar analysis was performed within individual subjects, for each of the three subjects that had a significant (corrected) high-frequency duration-tracking response, in the electrode where the effect was maximal (either O2 or PO8). Again, we found no differences between trials without or with high rate of saccades for any of the three subjects (all p-values >0.05, Figure 5B). We also verified that duration-tracking responses could be found in the absence of any saccades. Significant duration-tracking was found both for the group average ERP (Figure 5C) as well as for high frequency responses in individual subjects (Figure 5D) when including only trials with no saccades in the analysis, with a similar scalp topography as the results of the full analysis. This outcome is consistent with the results of a similar analysis of the intracranial sustained visual high-frequency response, showing the same insensitivity to saccades occurring during stimulus presentation (Gerber et al., 2017).

**Figure 5:**
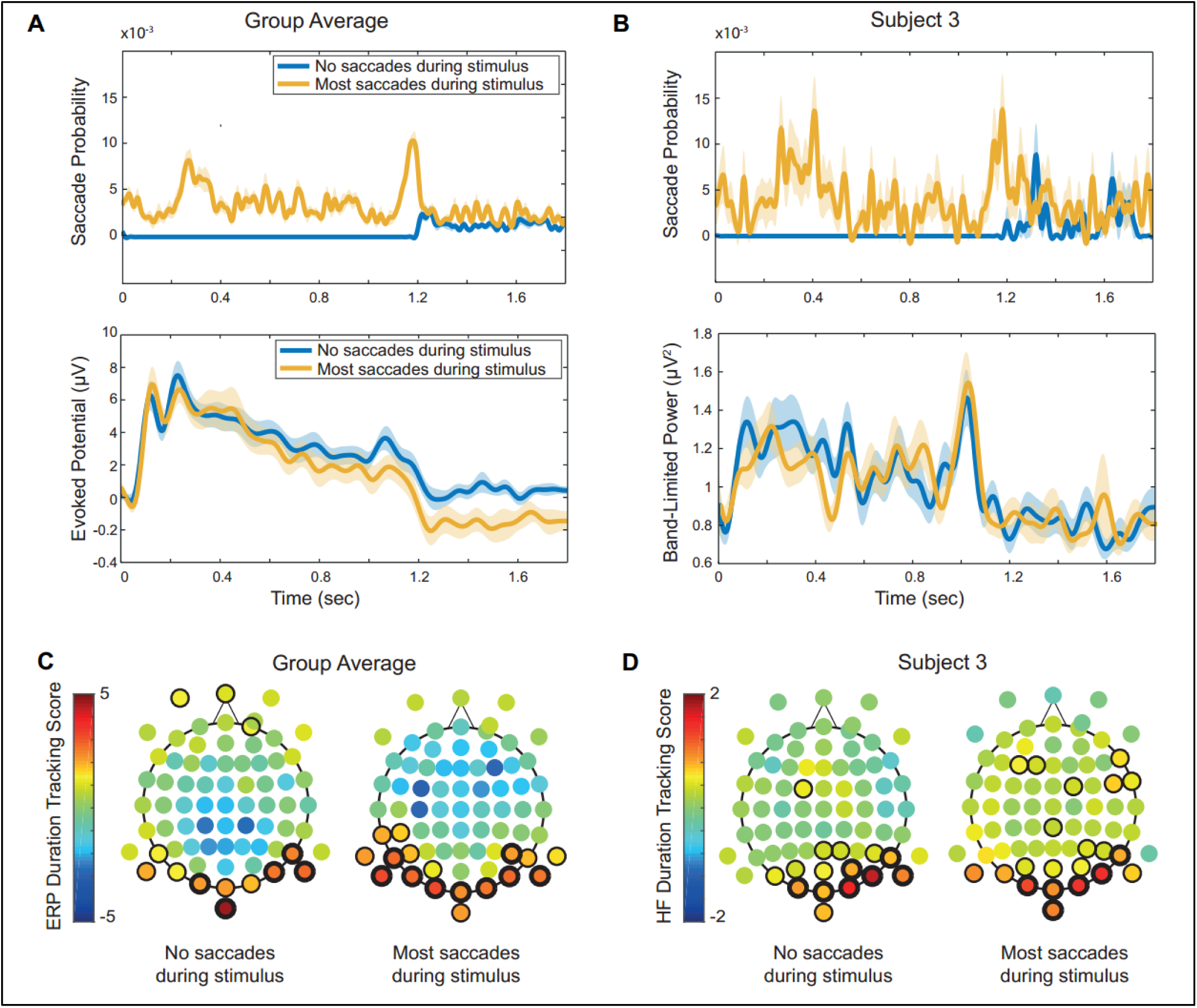
Experiment 1 – sustained baseline shift and high frequency responses are not driven by saccades. To test the hypothesis that the sustained baseline shift and high-frequency are the result of averaging transient responses driven by eye movements, each subject’s single trials were split into a group containing no saccades and a group containing the same number of trials with the highest number of saccades. **(A)** Top: Mean saccade rate, averaged across subjects, where in each subject trials were averaged separately for each saccade rate group (in the high saccade rate group there were 4.7 saccades per second on average across subjects). Bottom: Mean event related potential in electrode Iz, averaged across subjects, for the same trials analyzed in the top plot. No significant differences were found between the two averages. **(B)** Since high frequency duration-tracking responses cannot be seen in the group average, single subjects were analyzed individually for the effect of saccade rate on the sustained high frequency response. Top: Mean saccade rate for the two saccade rate groups in subject 3 (there were 5.1 saccades per second in the high saccade rate group for this subject). Bottom: The mean high frequency response in electrode O2, for the two trial groups averaged in the top plot. No significant differences between the averages were found in this subject or in any other subject with significant duration-tracking high frequency responses. **(C)** Duration-tracking analysis was carried out on the group-average ERP for each saccade rate group (left: trials with no saccade; right: trials with highest number of saccades. Thin outlines indicate p<0.05, thick lines indicate p<0.05/72). Significant duration-tracking responses were found in both conditions with a similar scalp distribution as for the undivided data, supporting the conclusion that sustained activity is not contingent on saccades. **(D)** Duration tracking analysis of the high frequency response in subject 3. Significant duration-tracking responses were found in both conditions in each of the analyzed subjects.

### Experiment 2

The sustained high-frequency and baseline shift responses found on the scalp are highly reminiscent of the two corresponding types of sustained responses we previously found, under similar experimental conditions, in intracranial recordings of early visual cortex (Gerber et al., 2017). If indeed the scalp-recorded response originates in early visual cortex, it should be strongly retinotopic – right and left stimuli should induce responses predominately in the contralateral hemisphere, and upper and lower visual fields should have different scalp topography and morphology. Specifically, the cruciform anatomy of early visual cortex in and around the calcarine sulcus suggests that evoked responses of these regions should present with opposing polarities on the scalp (Ales et al., 2010; Di Russo et al., 2002; Jeffreys, 1971; Kelly et al., 2013). We tested these predictions in a new experiment using stimulation in one of the four quadrants of the screen, excluding the center (Figure 1B). The stimuli were black and white checkerboard wedges presented for 300, 900 or 1500 ms, and the subjects’ task was to press a button whenever a word that was briefly flashed over the central fixation cross was a name of an animal (hit rate: mean 94%, std. deviation 5.2%; false alarms: mean 0.1%, std. deviation 0.1%). To increase statistical power, we defined a region of interest (ROI) based on the results of experiment 1 for high-frequency and baseline shift duration-tracking responses, consisting of 19 electrodes covering the posterior part of the scalp (Iz, Oz, O1-2, POz, PO3-4, PO7-8,P5-10, TP7-8, M1-2).

#### Duration-tracking in the event related potential

Surprisingly, the duration-tracking baseline shift was largely absent in the responses to the peripheral checkerboard wedges presented in the second experiment (Figure S2A). No electrode showed a significant duration-tracking baseline shift in the subject average for any of the four stimulus positions (after Bonferroni correction for the 19 posterior electrodes in the ROI). Analyzed individually in each subject, significant duration-tracking responses appear in only five electrodes across all subjects and stimulus conditions, with no consistent topography between conditions or subjects. This is in sharp contrast with the results of experiment 1, where the duration-tracking baseline shift was very robust in the subject average as well as appearing in nearly every individual subject’s recordings. This difference can be interpreted as a result of the difference in stimulus properties – specifically its position (foveal vs. peripheral), size, and content (complex meaningful images vs. simple shapes). As expected, stimulation in the upper vs. the lower visual field resulted in polarity inversion of the early evoked potential (C1 component, ∼80 ms) in posterior electrodes (Figure S2B). However, the absence of duration-tracking baseline shift precluded the analysis of polarity inversion in the sustained visual evoked potential.

#### Duration-tracking in the high frequency power

At the individual subject level, as in experiment 1, a subgroup of subjects exhibited a statistically significant duration-tracking response in the high frequency band. In those subjects the response was clearly lateralized, appearing in posterior electrodes contralateral to the stimulus position (Figure S3). We additionally observed that high frequency responses appeared exclusively for stimuli in the bottom and not upper visual field (i.e. only for lower-left and lower-right stimuli). At the group level, a significant duration-tracking response was found in two right posterior electrodes (O2 and PO4) in the lower-left stimulus condition, whereas the duration-tracking response in left-posterior electrodes for lower-right stimuli was also evident though non-significant after Bonferroni correction (Figure 6). The strong lateralization as well as sensitivity to altitude of the high-frequency response is consistent with the hypothesis that its source is found in early visual cortex. In comparison to experiment 1, the high frequency responses seen in experiment 2 appear to have a narrower bandwidth (i.e. approx. 60-80Hz as opposed to 50-100Hz, compare Figure 6 and Figure S3 to Figure 3. Also see similar narrowband responses in experiment 4, Figure 11). This difference is consistent with the finding that intracranial high frequency visual responses are narrower for high contrast grating stimuli than for grayscale noise patterns (Hermes et al., 2014).

**Figure 6:**
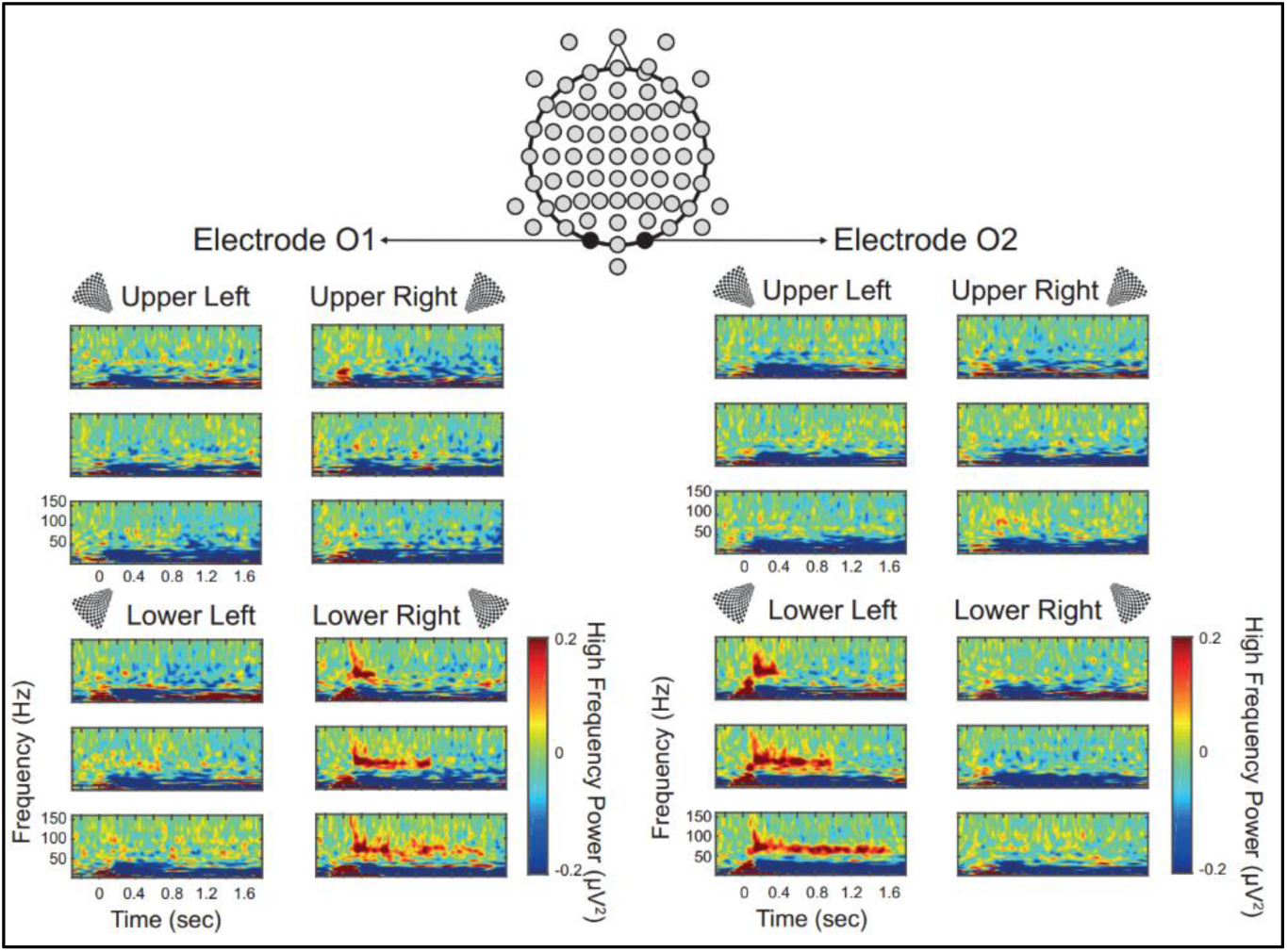
Experiment 2 - high frequency sustained responses to peripheral checkerboard stimuli. Group average time-frequency plots for checkerboard wedge stimuli presented in four peripheral locations in experiment 2, in two posterior electrodes. Image on top shows the positions of electrodes O1 and O2 on the scalp. For each electrode and stimulus location, three plots show the group average time-frequency response for the 300, 900, and 1500 ms stimulus duration from top to bottom. A clear lateralized response can be seen with lower left stimuli eliciting a response in right posterior electrode O2 (duration-tracking score 0.4, p < 0.005), and lower right stimuli eliciting a response in the left posterior electrode O1 (duration-tracking score 0.07, p < 0.05).

### Experiment 3

The differences between the results of experiment 1 and experiment 2 suggest that the visual duration-tracking baseline shift recorded on the scalp is strongly dependent on stimulus properties. To investigate this dependence directly, experiment 3 was designed to address two aspects in which Experiments 1 and 2 differed: stimulus eccentricity (the stimuli in experiment 1 included the fovea while in experiment 2 the stimuli avoided the center of the visual field), and semantic content (the stimuli in Experiment 1 were meaningful, whereas in Experiment 2 they had low semantic content). Stimuli in Experiment 3 were either black and white checkerboard rings with a small (more central), intermediate, or large (peripheral) eccentricity, or central disks with a black and white “Mooney” image that depicted a face or object, or a scrambled meaningless image (Fig. 1). Mooney images were used rather than natural images so that the low level features of the semantic categories will be more similar. To reduce the duration of the experiment, only two different stimulus durations were used in this experiment (500 and 1000 ms). To reduce confounding stimulus eccentricity with the size of the activated cortical area, the size of the ring stimuli was normalized according to the cortical magnification factor at each eccentricity (see Materials and Methods). Subjects’ task was to respond with a button press to a subset of scrambled Mooney images that were arranged symmetrically (hit rate: mean 94.5%, std. deviation 4.5%; false alarms: mean 1.5%, std. deviation 3.2%).

#### Duration-tracking in the event related potential

The checkerboard ring stimuli and image stimuli were analyzed separately. To define a region-of-interest for the comparison between the 3 ring sizes and between the different Mooney images, duration-tracking analysis was first performed on the average of the three ring conditions, and separately on the average of the three Mooney image conditions. For the average of the ring stimuli, the maximal duration-tracking effect size was found in midline occipital electrode Iz (duration-tracking score 0.83, p = 0.0095, not significant after Bonferroni correction for 19 electrodes). This electrode was therefore used for subsequent comparison between stimulus eccentricity conditions. For Mooney images, a significant duration-tracking effect was found in five posterior electrodes with a maximum at left parieto-occipital electrode PO7 (duration-tracking score 3.32, p < 10^−3^), which was used for subsequent analysis (Figure 7A). Then for each condition and subject, we took the average potential relative to the baseline within a 0.75 to 1.05 sec post-stimulus window, which excludes as much as possible any onset- or offset-driven response components (Figure 7B). This parameter was used as dependent variable to compare between conditions across subjects, using repeated-measures one-way ANOVA.

**Figure 7:**
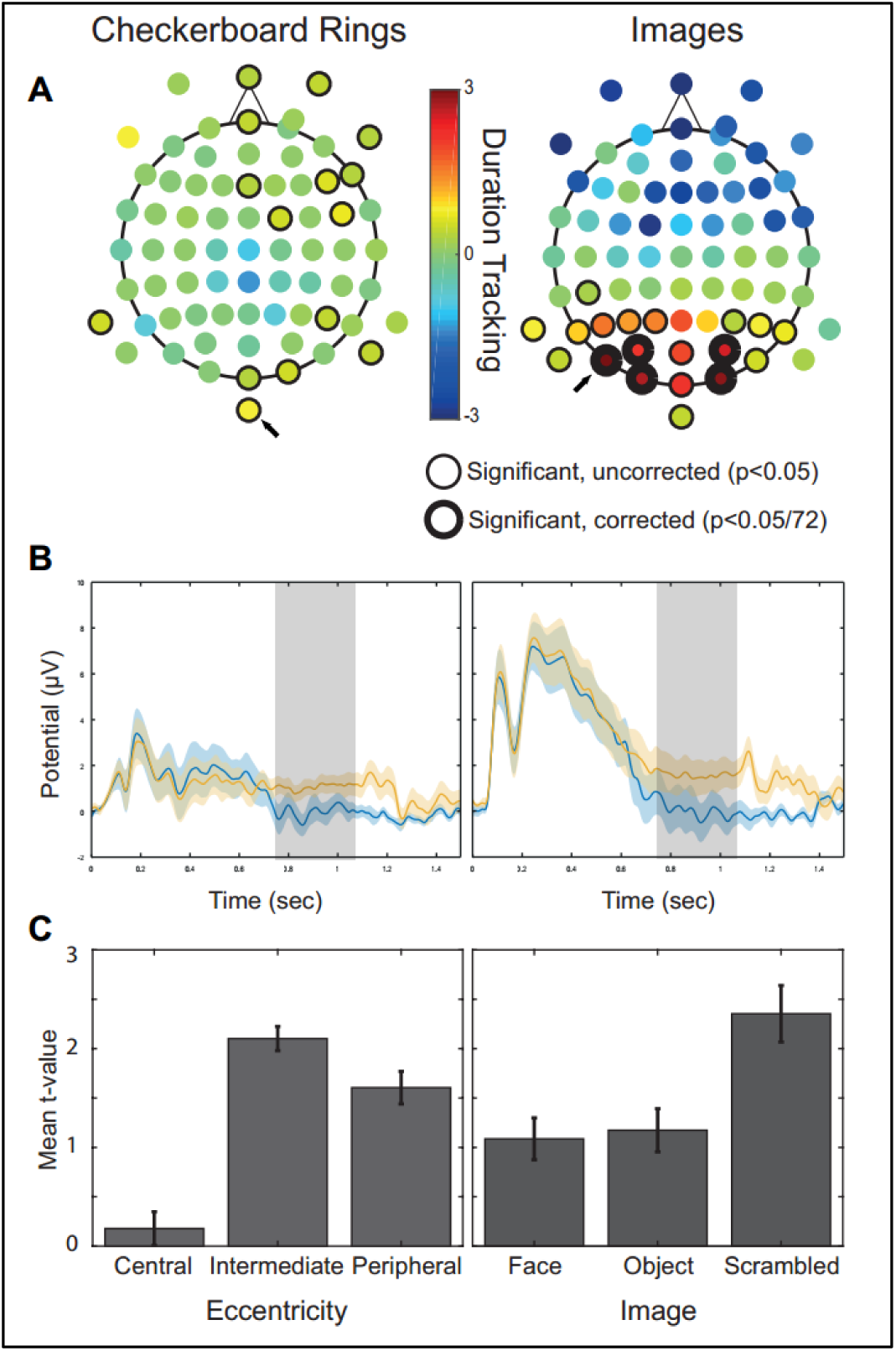
Experiment 3 – comparison of the sustained visual baseline shift response between stimulus conditions. The magnitude of the sustained visual baseline shift in experiment 3 was compared among three stimulus eccentricity conditions (checkerboard rings with small, intermediate and large eccentricity) and three semantic content conditions (central face, object, or scrambled image). For each comparison, an electrode region-of-interest was selected as the electrode with largest duration-tracking effect after averaging over the three conditions. **(A)** Topography of the duration-tracking effect for each three-condition average. Left: checkerboard ring stimuli; the largest effect size was in electrode Iz (marked with arrow). Right: central image stimuli; the largest effect size was in electrode PO7 (marked with arrow). **(B)** Group-average responses in the selected electrodes, averaged across the three respective conditions for each comparison. Note that the plots do not necessarily reflect the differences in duration-tracking score between the checkerboard and image condition groups as shown in (A), since this measure is dependent on the variance of within-subject differences that is not seen in the group average. Gray area indicates the time window for subsequent analysis of differences between conditions. **(C)** ANOVA results for each condition comparison, based on the mean t-value of the difference between the short- and long-duration responses within the analyzed window in each condition. A significant difference was found between the small and intermediate eccentricity checkerboard rings.

For checkerboard ring stimuli, we found a significant difference between conditions (F(2,24) = 4.54, p<0.05). There was a larger response for the intermediate ring than for the central ring (t(12)= 2.98, p<0.05), and no significant difference between the intermediate and peripheral ring conditions (Figure 7C). For Mooney images, ANOVA did not reveal any significant difference between conditions. Analyzed separately, a significant (corrected) duration-tracking effect was found in posterior electrodes for the peripheral ring, face image, and scrambled image conditions, with a non-significant but typical posterior effect topography in the intermediate ring and object image conditions (Figure S4). The larger response to the intermediate and peripheral compared to the central stimulus was unexpected given the robust duration-tracking responses observed for the central stimuli in experiment 1 but not for to the peripheral stimuli in experiment 2. This could be due to over-compensation for differences in cortical magnification, which led to the small eccentricity stimulus being much smaller than the larger-eccentricity stimuli. The compensation for cortical magnification was derived from data on cortical magnification in V1, while it is not clear whether the source of the sustained baseline shift in indeed in this area or restricted to it. Overall, the results suggest that sufficiently large stimuli produce a sustained visual baseline shift. The results from the Mooney images provides no evidence of an effect of semantic content.

#### Duration-tracking in the high frequency power

As in the previous experiments, we found significant high frequency duration tracking in posterior electrodes in a minority of subjects. Of the 14 subjects, one showed a single-electrode significant effect in a single condition, and three had a significant effect in at least one posterior electrode in either 3, 4, or 5 of the conditions (but never in the central ring condition, Figure S5). Where the effect was found, it tended to concentrate consistently around midline posterior electrodes. We did not carry out a systematic comparison between conditions because no adequate ROI for this comparison could be defined (the paucity of subjects showing duration-tracking precluded group analysis, and within subjects the topography of significant duration-tracking was not consistent enough between conditions to select an ROI that would not be biased toward a particular condition). Nevertheless, the absence of significant effects in the small eccentricity condition for both the low and high frequency signals suggests that this stimulus was insufficient to produce a robust sustained response of either type.

### Experiment 4

The results of experiment 3 indicated that the absence of sustained visual baseline shift in experiment 2 was likely due to the stimuli being insufficiently large for peripheral presentation. The goal of experiment 4 was therefore to revisit the question of lateralization and polarity inversion in the sustained visual baseline shift, which was left unanswered in experiment 2, using larger visual stimuli. In the high-frequency domain, it was established in experiment 2 that sustained broadband activity is strongly lateralized, indicating an early visual cortex source. We now sought to investigate whether high frequency sustained activity in high-level visual cortex could be detected on the scalp, reflected as differences in the topography of the high-frequency sustained response between simple and complex stimuli (grating vs. face images).

Stimuli were presented in a rectangular area covering nearly half of the screen, on either the top, bottom, left or right side of a central fixation cross, and were either a semi-circle of black and white grating or a grayscale frontal photo of a face. The duration of the stimuli was either 300 or 900 ms. Subjects were required to attend to each stimulus without moving their ayes from the fixation cross, and respond with a button press to target stimuli which faded gradually over the last 200 ms of their presentation rather than abruptly disappearing. The subjects performed well in this task, indicating that they have paid attention to the stimuli (hit rate: mean 88.9%, std. deviation 6.4%; false alarms: mean 0.8%, std. deviation 0.6%). In order to elicit polarity inversion in the evoked response, the stimuli had to activate the top or bottom visual field with minimal overlap. Therefore, trials where the average gaze position was more than 0.2 visual degrees toward the stimulus position (corresponding to the border of the stimulus itself, see Materials and Methods) were excluded from analysis. This strict requirement, in addition to exclusion of trials with amplitude artifacts in the unfiltered or high-frequency signal, led to a low number of trials remaining for analysis in some of the subjects. Any subject with less than 25% of total trials remaining for analysis were excluded, leading to 10 of 30 subjects being excluded from the analysis of both ERP and high-frequency responses.

#### Duration-tracking in the event related potential

Overall, increasing the stimulus size relative to experiment 2 had the desired effect of eliciting the duration-tracking baseline shift in the group average. Averaging the data from both face and grating stimuli, significant duration-tracking was found at the group level in 1-4 posterior electrodes for the four stimulus positions (Figure 8A). Similar results were found when the face and grating stimulus conditions were analyzed separately.

**Figure 8:**
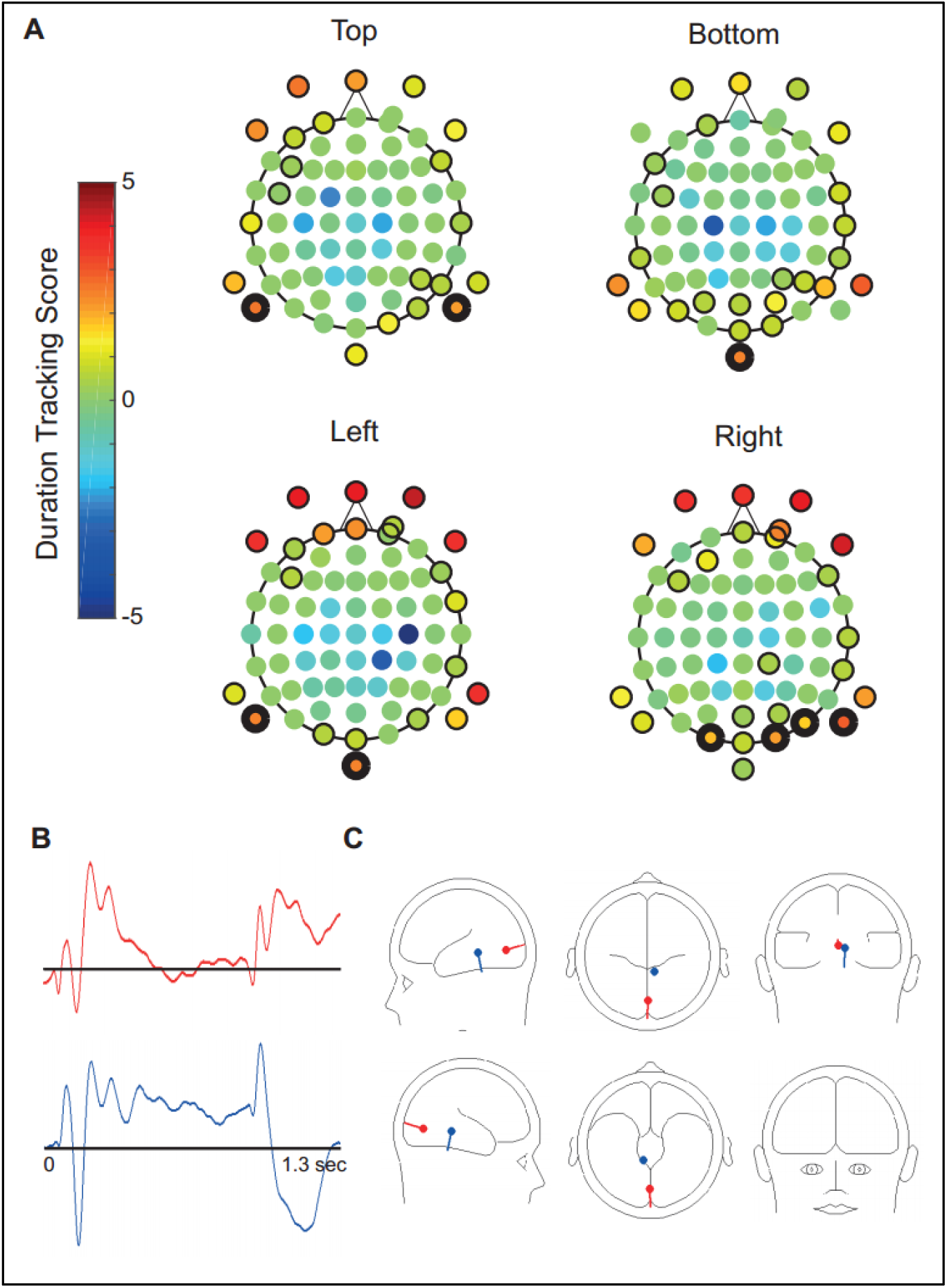
Experiment 4 – topography and diploe-source estimation of the sustained visual baseline shift. **(A)** Example stimulus and topography of the ERP duration-tracking score for each stimulus position, averaged across both face and grating stimuli. Thin outline indicates p<0.05 significance level and thick outline indicates significance after Bonferroni correction for the 19 posterior region-of-interest electrodes. Robust duration-tracking responses appeared in posterior electrodes but also in temporal and facial electrodes. **(B)** Topography of the duration-tracking score, for the average responses across all stimulus conditions. **(C)** To check whether this effect topography could correspond to any plausible cortical source configuration, we applied dipole source analysis to the average response across all stimulus conditions. 71% of the variance could be explained with two dipole sources, with one containing the sustained response component. **(D)** Coordinates of the two dipole sources shown in (C).

A striking feature of the duration-tracking baseline shift in this experiment was that it did not appear to have an occipitally-centered topography; rather, sustained responses were found most strongly in posterior electrodes, but were also seen in frontal (including facial) and lateral electrodes for all stimulus positions (Figure 8A,B. Note that frontal channels are not marked as significant after correction in Figure 8A since they were not within the pre-defined ROI). Whereas this pattern is most conspicuous in the results of Experiment 4, we observed that in the previous experiment (with the exception of the response to the Mooney images in Experiment 3) the minimum of the sustained effect was similarly in central scalp electrodes (compare to Figure 3C). To explore the possible source configuration that would generate this type of distribution, we applied dipole modeling to the data using the BESA tool (BESA 6.0, http://www.besa.de). The BESA software uses a four-shell ellipsoidal head model and searches within the model for the position and orientation of one or more electrical source generators that will explain the maximal amount of variance for the observed data on the scalp (Berg and Scherg, 1994; Scherg and Picton, 1991). Since the sustained response topography appeared to be roughly similar for all stimulus conditions (i.e. type and position), we fitted a dipole source model to the average response to 900 ms stimuli, across all subjects and stimulus conditions, over a time window of 1.4 seconds. The analysis showed that 71% of the variance in the observed activity on the scalp could be explained by a midline occipital source with onset and offset components, and an additional vertically-oriented source more anteriorly, containing the sustained response component (Figure 8C,D). Adding additional dipoles to the model increased the explained variance but did not add sources with a clear sustained component. The same vertically oriented central dipole appeared consistently when applying the model separately to the top, bottom, left or right stimulus conditions, or to the isolated sustained response component derived as the difference between the response to the 900 ms and 300 ms stimulus durations. We interpret the results of this analysis not as indicating a specific source location for the sustained response (as the dipole localization is too crude), but as confirming that a cortical source configuration exists that could produce the observed scalp topography, and suggesting that the source of the sustained response is likely different from the source of the evoked onset and offset response.

#### Polarity inversion and lateralization of the sustained baseline shift

We compared the evoked response to stimuli in the top vs. the bottom visual field to test for polarity inversion of the sustained response. Polarity inversion is typical of visual responses arising from early visual cortex where cortical dipoles in the superior and inferior banks of the calcarine sulcus are oriented in opposite directions in relation to the scalp, thus manifesting as opposite-polarity deflections in the EEG. Looking at early visual evoked responses, we found the expected polarity inversion for upper vs. lower visual field stimuli of the C1 component (∼80 ms peak latency) in mid-line posterior electrodes, with a peak at electrode POz, as described in the literature (Di Russo et al., 2002) (Figure 9A). At the same time, the sustained response component was consistently positive and did not differ between upper and lower visual field stimuli at any electrode (t-test of the mean potential in the 600-900 ms window for 900 ms duration trials in top vs bottom stimulus trials, variance across subjects, Bonferroni corrected for 72 electrodes, Figure 9B). The lack of polarity inversion in the sustained response argues against the assumption that its source is in V1 (although this conclusion should be taken with caution as V1 sources may not necessarily produce polarity inversion (Ales et al., 2010)).

**Figure 9:**
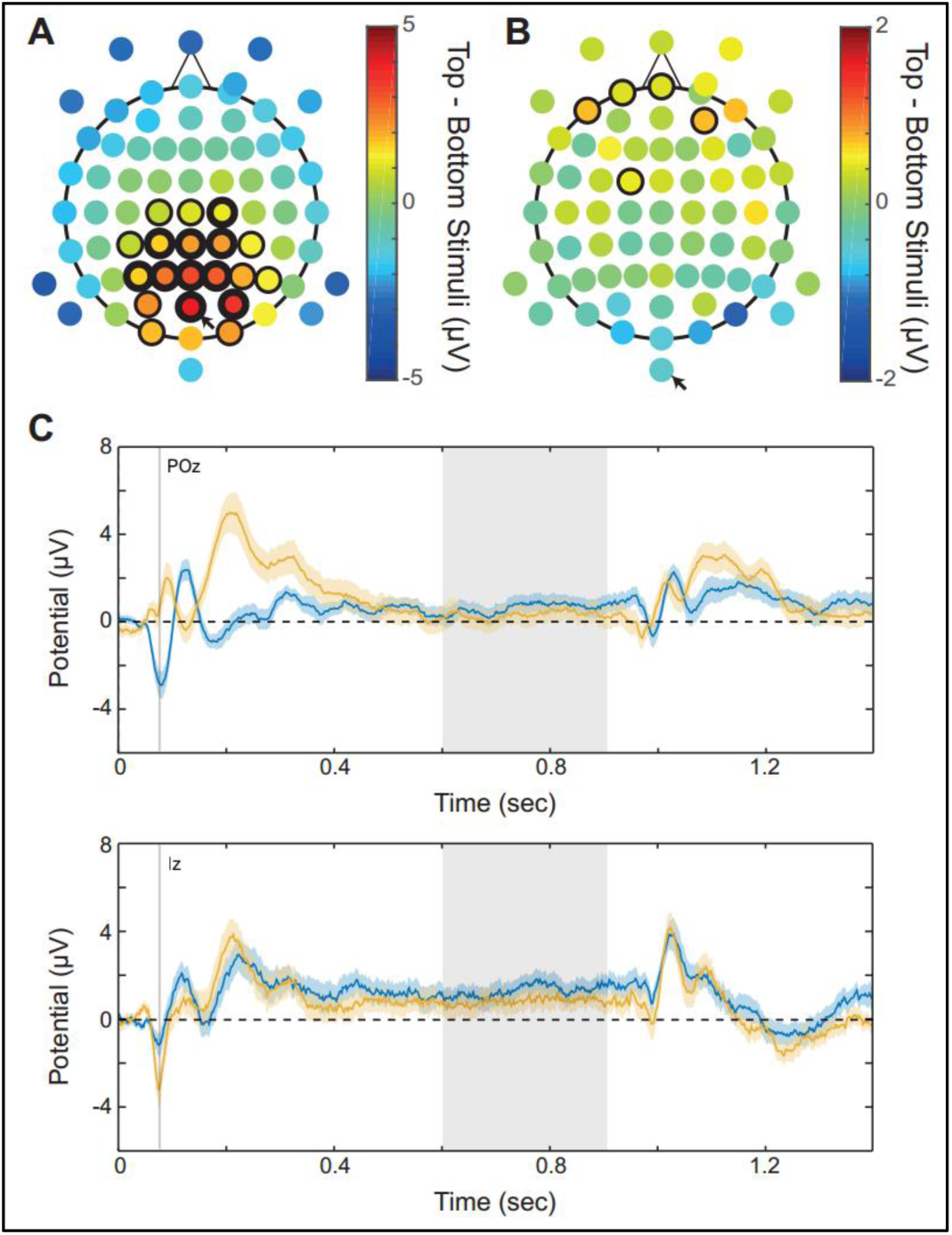
Experiment 4 – polarity inversion in the sustained visual baseline shift. **(A)** Topography of the group-mean difference between the top and bottom visual field stimuli at the 80 ms latency (corresponding to the C1 component). As predicted by the literature, a polarity inversion effect can be seen in posterior-parietal sites with a maximum in electrode POz (marked with arrow). Thin and thick outlines correspond to t-test significance before and after Bonferroni correction for 72 electrodes respectively. **(B)** Topography of the group-mean differences between the top and bottom visual field stimuli, averaged across a 600-900 ms latency window (corresponding to the sustained response component). A polarity inversion effect similar to that found for C1 is not found for the sustained response. **(C)** Group-average responses for top and bottom visual field stimuli. Top: electrode POz, showing polarity inversion in C1 but not in the sustained response. Bottom: electrode Iz, where the sustained response is most robust, also does not show polarity inversion of this component. The gray patches indicate the latency of the C1 and the sustained response window.

An additional hypothesis was that a retinotopic source of the sustained baseline shift would produce lateralization of the response for stimuli presented in the left vs. right visual field. We compared across subjects the magnitude of sustained responses, measured as the mean of the 600-900 ms window in 900 ms duration trials, in left vs. right visual field stimuli. While no single electrode passed the significance threshold after Bonferroni correction for 72 electrodes, the resulting topography suggests a moderate lateralization of the sustained response component, with ipsilateral, rather than contralateral electrodes exhibiting stronger sustained activity (Figure 10).

**Figure 10:**
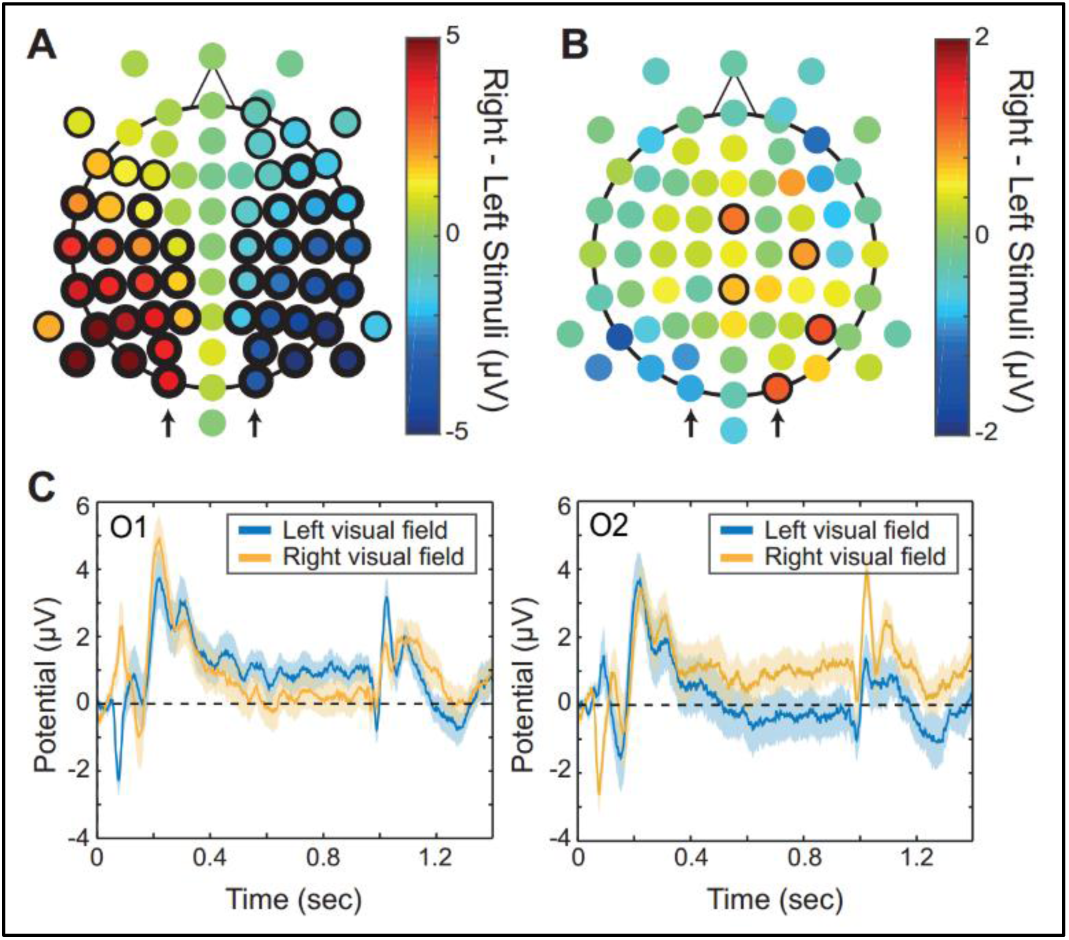
Experiment 4 – lateralization in the sustained visual baseline shift. **(A)** Topography of the group-mean difference between the right and left visual field stimuli at the 80 ms latency. Thin and thick outlines correspond to t-test significance before and after Bonferroni correction for 72 electrodes respectively. **(B)** Topography of the group-mean difference between right and left visual field stimuli, averaged across a 600-900 ms latency window (corresponding to the sustained response component). While no single electrode shows a significant difference after Bonferroni correction, the sustained response appears to be lateralized with stronger sustained activity in electrodes ipsilateral to the stimulus. **(C)** Group-average responses for 900 ms stimuli in the left and right visual field, for left and right posterior electrodes O1 and O2, marked with arrows in (A) and (B).

#### Duration-tracking in the high frequency power

The high frequency responses to grating stimuli were similar to those observed in experiment 2 for peripheral checkerboard stimuli. In the group average, significant duration-tracking high frequency responses were found in lateral posterior electrodes contralateral to the left/right visual field stimulus, and in a right posterior electrode for the bottom visual field stimulus (Figure 11A,B). As in experiment 2, a significant response was observed for stimuli in the lower but not upper visual field. Face stimuli were also analyzed in an attempt to observe differences in response topography attributable to activity in high-level, face-selective visual areas; however, the lateralized faces failed to elicit sufficiently robust high-frequency sustained responses in this group of subjects (Figure 11C).

**Figure 11:**
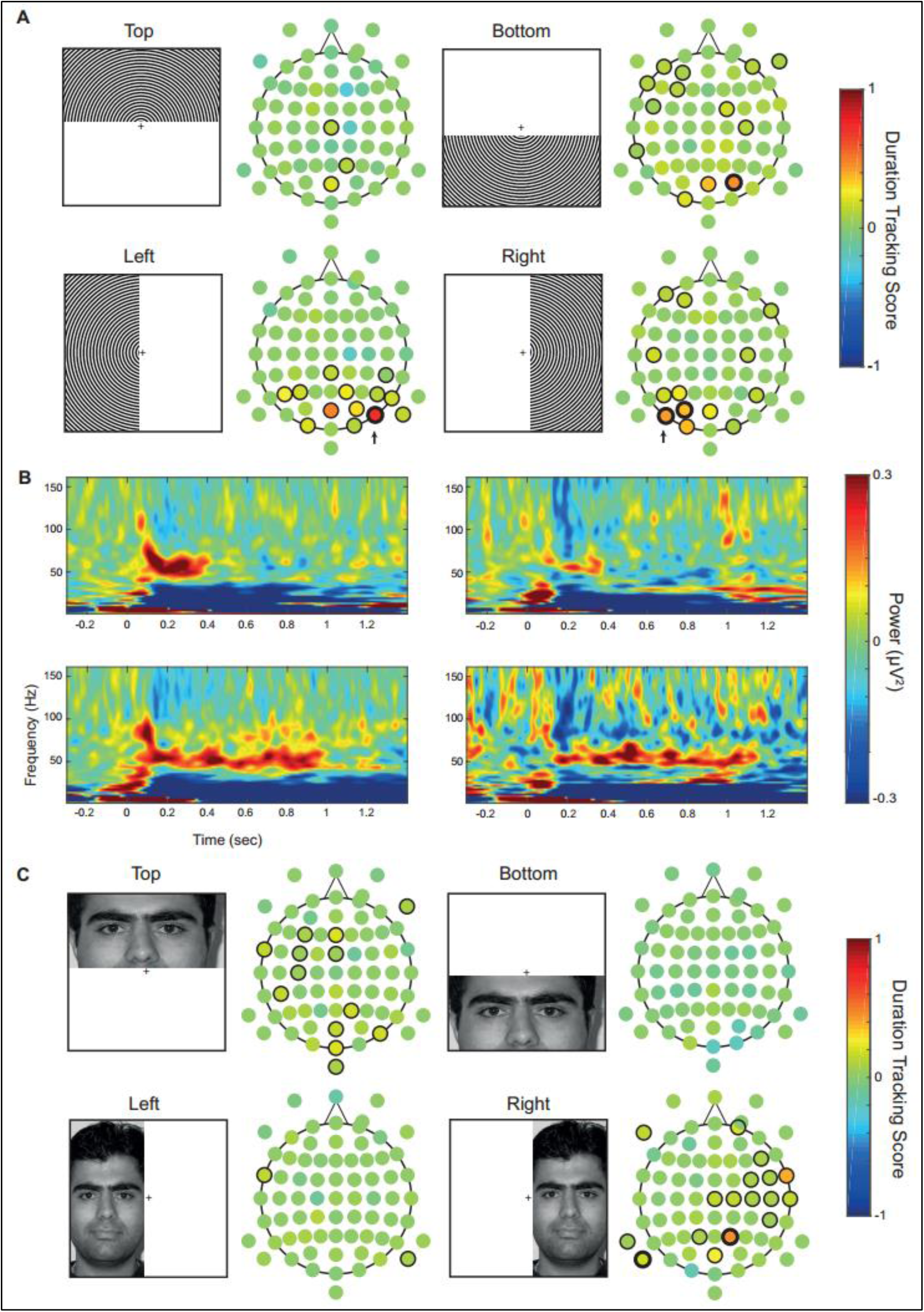
Experiment 4 – high frequency sustained responses for peripheral face and grating stimuli. **(A)** Example stimulus and topography of the high-frequency duration-tracking score for each stimulus position in the grating image condition. Thin outline indicates p<0.05 significance level and thick outline indicates significance after Bonferroni correction for the 19 posterior region-of-interest electrodes. Similarly to the results of experiment 2, we found a lateralized duration-tracking high frequency response that was absent for upper visual field stimuli. **(B)** Group-average time-frequency plots for electrode PO7 (left) and right visual field stimuli and electrode PO8 (right) and left visual field stimuli. These electrodes are marked with arrows in (A). Top and bottom plots correspond to 300 and 900 ms stimuli. **(C)** Example stimulus and topography of the high-frequency duration-tracking score for each stimulus position in the face image condition. High-frequency responses were generally absent in this condition.

## Discussion

What kind of neural activity is associated with perceiving a static visual stimulus beyond the initial moment of its appearance? We have previously addressed this question using intracranial recordings in humans and found that high-frequency broadband power responses were sustained throughout stimulus presentation in the early visual cortex, with diminishing moment-to-moment reliability downstream the ventral visual pathway. In a small subset of EVC sites we have also found sustained positive or negative potential shifts lasting for the entire duration of the stimulus. These responses were not correlated with the high frequency broadband responses – some sites showed both phenomena while others showed only one. In the current study, we implemented a similar variable-duration stimulation paradigm to explore what kind of duration-tracking activity is observable in scalp EEG recordings.

The results of our first experiment showed that both phenomena, namely the sustained baseline shift and sustained high-frequency response, can be recorded on the scalp in response to foveal images of objects and faces. The sustained baseline shift had a widespread topography with a posterior maximum and was detected in nearly all individual subjects.

While a sustained visual baseline shift with roughly similar morphology has been demonstrated before (Maquet et al., 1996; N’Diaye et al., 2004; Pouthas et al., 2000), it was to our knowledge only in the context of explicit duration estimation tasks, whereas in the present case the duration of the stimuli was irrelevant to the task. We therefore demonstrate that the sustained duration-tracking response is a task-unrelated, mandatory component of the electrophysiological response to visual stimulation. The sustained broadband high-frequency response had a highly localized right-posterior topography. Whereas at the single subject level it was evident only in a subset of the subjects, the pattern of the responses in these subjects, the precise correspondence with the duration of the stimuli, and the striking resemblance to our previous intracranial recordings, provide unequivocal evidence that broadband high-frequency responses can, in some cases, be recorded on the scalp. This may have particular importance as intracranial studies of the last few years have established this type of response as a reliable proxy for activation of populations of neurons, correlated with multiunit firing rates (Manning et al., 2009; Miller et al., 2014, 2009; Ray and Maunsell, 2011). That said, consistently with previous findings from scalp EEG (Keren et al., 2010; Muthukumaraswamy et al., 2010; Whitham et al., 2007), the signal-to-noise ratio of this signal was too low to detect significant post-stimulus power modulation in many subjects, many of whom did not manifest any kind of visual response in the high-frequency range. This variability between subjects can potentially be attributed to noise from muscle activity and external sources, which was not sufficiently suppressed by our pre-processing, but also to anatomical differences in the mapping of activation to specific cortical patches, and how these patches are oriented relative to the recording electrodes. The latter possibility is commensurate with the highly local effects found for the broadband high frequency effect in the current study (e.g. 1-2 posterior electrodes). Variability in skull shape or thickness may also play a role.

We note the striking difference in how low and high frequency neural responses translate from intracranial recordings to scalp recordings: sustained high frequency responses were relatively ubiquitous in the visual cortex measured intracranially, but appeared only in some subjects with a very localized scalp topography using scalp EEG, whereas the sustained visual baseline shift appeared over a wide posterior scalp topography and in nearly every EEG subject despite being found in only few nearby sites on the early visual cortex in the intracranial dataset. This result suggests that the projection of different neural signals from brain to scalp may serve not only to attenuate some neural signals, but also to make other cortical sources of activity more accessible.

Methodologically, the finding that high frequency visual responses on the scalp are sustained throughout visual stimulus presentation can help isolate neural from artifactual sources of high frequency activity in relation to visual perception. Non-neural noise, both biological (muscle activity in the scalp and eyes, heartbeat, etc.) and electrical (power line noise, monitor refresh rate noise, etc.) may interfere with and even be erroneously interpreted as neural activity, especially if it is modulated by experimental manipulations (Yuval-Greenberg et al., 2008). However, a noise source that is modulated by the appearance of a visual stimulus is less likely to be sustained for the duration of the stimulus. Thus, “tagging” a high frequency response with a specific stimulus duration and observing a corresponding duration-tracking response may serve as an indication for the neural origin of the observed signal.

In experiments 2-4, we attempted to characterize the properties of the two types of sustained visual activity along two dimensions – retinotopic location and semantic content. When using peripheral stimuli, the high frequency sustained response showed a highly localized topography that appeared contralaterally from the stimulated visual field, indicating a retinotopic cortical origin. The fact that the high frequency response was hardly recorded for stimuli in the upper visual field probably stems from morphological factors, i.e. differences in how sources in inferior and superior parts of the early visual cortex are projected to the scalp, combined with the inevitable poor coverage of inferior part of the head (intracranially, sustained high frequency responses were found in both inferior and superior banks of the calcarine sulcus). The observation in the intracranial study that sustained high frequency visual responses are strongest in early visual cortex, in conjunction with the lateralization and focal topography of the scalp EEG high frequency response, suggest that the early visual cortex is the primary source of the EEG signal as well.

The source of the sustained baseline shift is less clear. Unlike the high frequency broadband response, the baseline shift had a broad scalp topography, with maximal amplitude found either in occipital electrodes or in frontal or temporal sites. Equivalent-Dipole modelling suggests that this topography is consistent with a source which was more anterior relative to the occipital pole than the separate dipole that explained the early evoked potential. Equivalent dipoles should not be mistaken for source localization as they represent a center of mass of activity, leaving open the exact location of the sources. We view this analysis as showing that the sources of the sustained responses are likely separable from those of the early evoked potentials. This is also consistent with the preservation of positive polarity of the baseline shift for upper and lower visual field stimulation, in contrast with the reversal of polarity of the C1 at 80ms. Nevertheless, it is still likely that the source of the sustained baseline shift is in the early visual cortex considering its strong dependence on low-level features of the stimulus (i.e. eccentricity and/or size).

Our study is the first step towards understanding sustained responses in EEG, bridging a gap towards intracranial recordings. We were able to replicate the two main sustained responses we previously found on the surface of the visual cortex. As we found intracranially, sustained baseline shift and high frequency responses in scalp EEG appear to be two distinct markers of sustained visual processing in the cortex, possibly corresponding to different intracortical sources and functional aspects of visual perception. The two sustained response types have different scalp topography and do not necessarily co-occur in the same electrode or even the same individual subjects. Our results also suggest different dependencies on stimulus properties for each response type. The variability of the duration-tracking baseline shift between experiment 1 and 2 and between the conditions of experiment 3 suggests that it is strongly dependent on the stimulus position and size in the visual field. The differences in the high frequency sustained response to grating and face stimuli in experiment 4 hint that it is strongly modulated by stimulus contrast and/or spatial frequency. Future studies are needed to shed more light on the specific characteristics of each response type and on the associated cortical processes.

## Supplementary Figures

**Figure S1:**
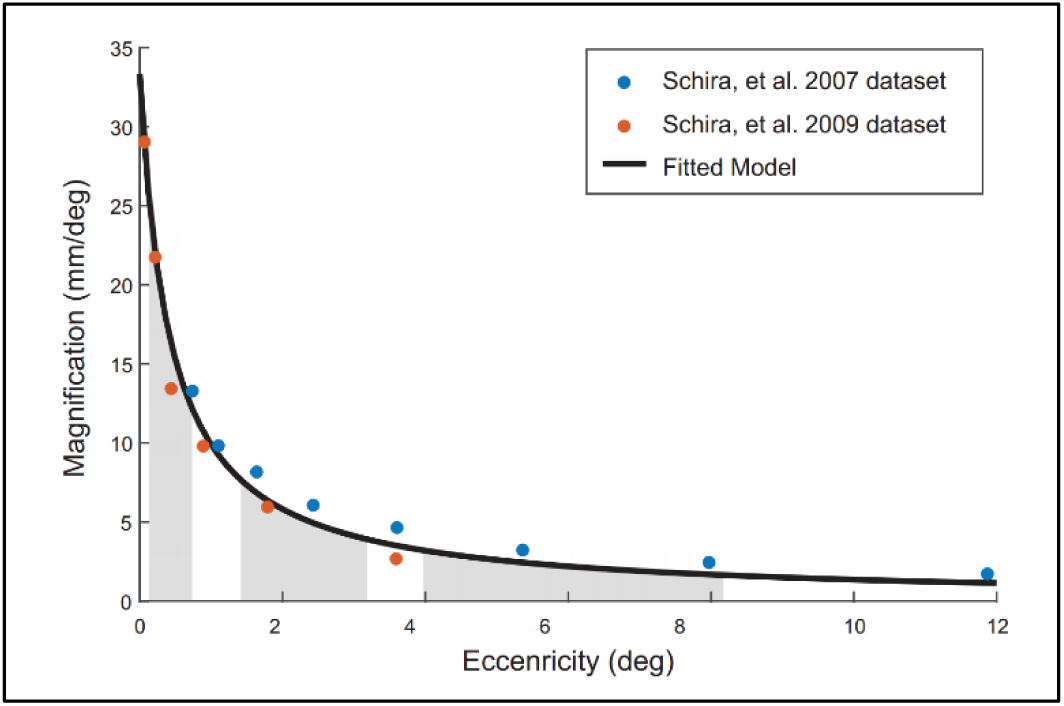
Experiment 3 – derivation of ring stimulus sizes with equal cortical area activation for different visual eccentricities. To generate ring stimuli with different eccentricities that are normalized in respect to activated cortical area, we first acquired cortical magnification data for the relevant eccentricity range (approx. 0.1 to 8 visual degrees). This data was extracted from published figures in (Schira et al., 2007), covering the 0.75-12 degree range and in (Schira et al., 2009) covering the 0-4.8 degree range for visual area V1. The two data sets were combined and a standard cortical magnification function *M*(*E*) = *k*/*E* + *a*, where *E* is eccentricity and *k* and *a* are fitted parameters. The figure shows the two data sets and the fitted function. The resulting fitted parameters were *k=19.2*, *a=0.77*. To equalize cortical magnification for two eccentricity ranges (i.e. from the inner to the outer ring border), we derived a formula based on equalizing the definite integrals of the function for these eccentricity ranges (gray patches in the figure). Solving the equality produces the formula (*y*_1_ + *a*)/(*x*_1_ + *a*) = (*y*2 + *a*)/(*x*2 + *a*), where *x_i_* and *y_i_* are the inner and outer eccentricity of ring *i*, respectively (parameter *k* is canceled out in this derivation). Using this formula, small, intermediate and large eccentricity rings were generated with equal estimated cortical activation areas for V1.

**Figure S2:**
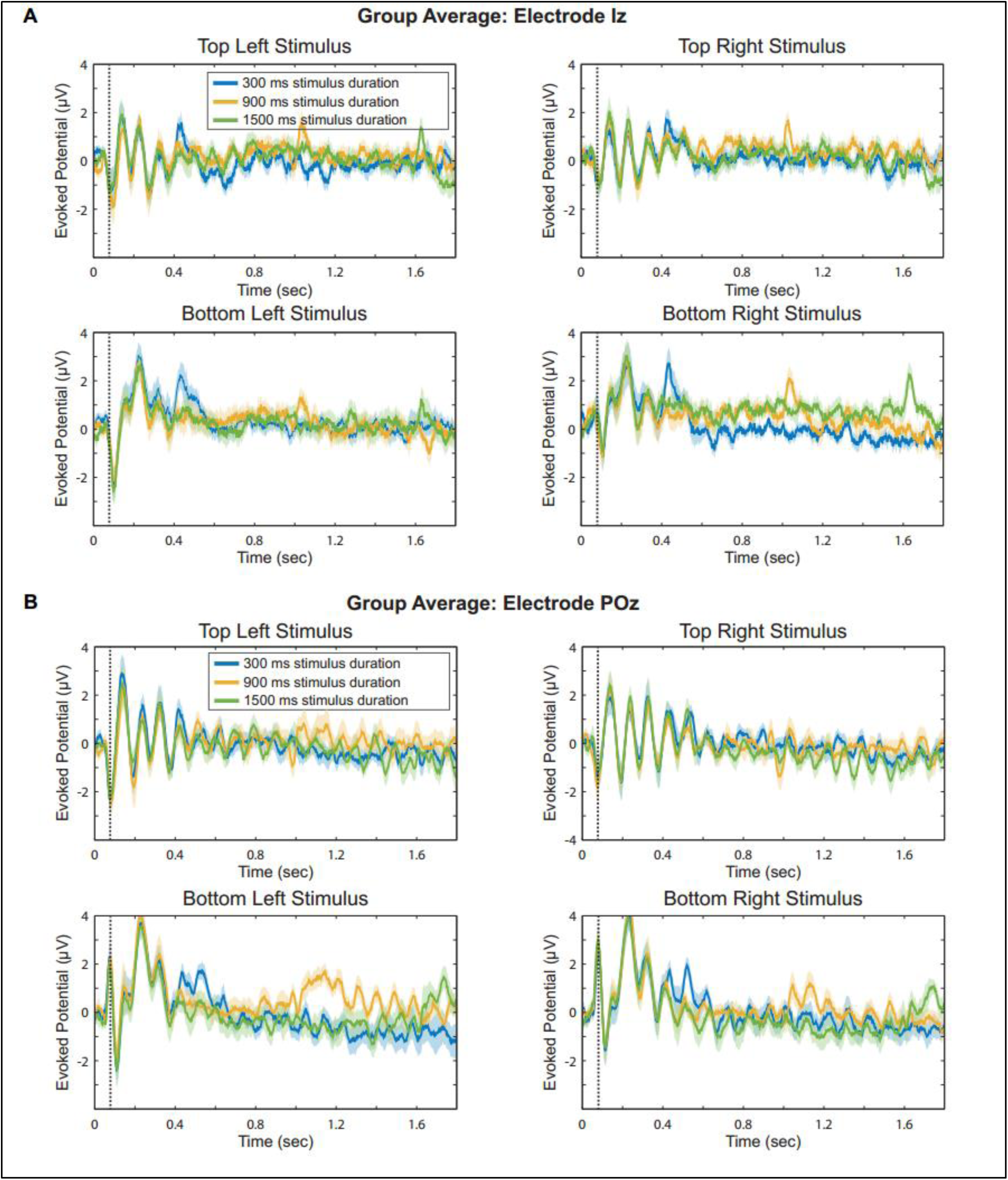
Experiment 2 – Polarity inversion of early evoked potentials but no sustained baseline shift. **(A)** Group average ERPs for the three stimulus duration in each of the stimulus position conditions, in electrode Iz, which showed the largest duration-tracking effect in Experiment 1. No ERP duration-tracking effect can be seen in this or any other electrode in Experiment 2. **(B)** Group average ERPs for electrode POz. In this electrode, the polarity inversion of the C1 evoked component can be seen between top and bottom stimulus positions (∼80 ms latency, marked with dotted line in each plot).

**Figure S3:**
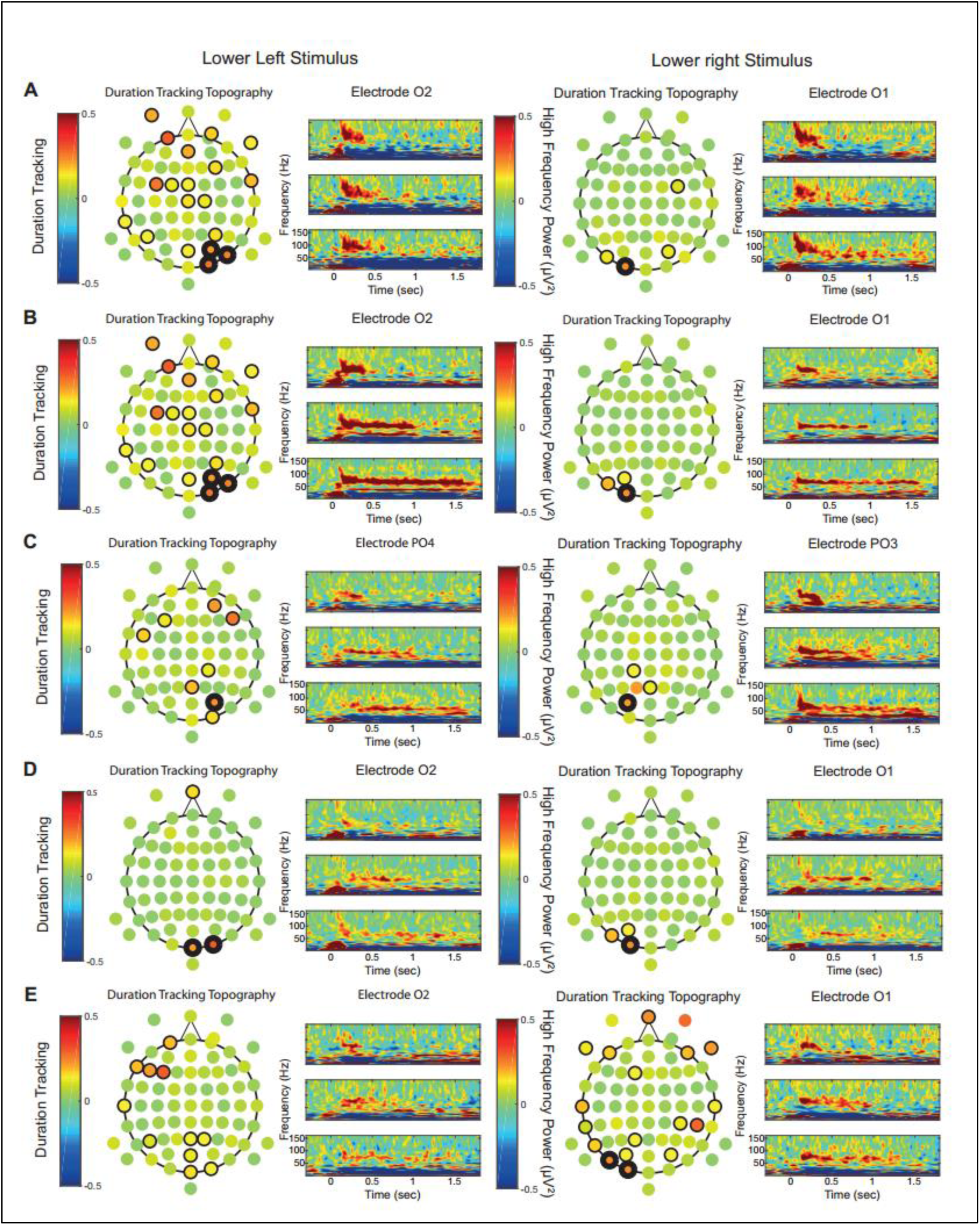
Experiment 2 – lateralized high frequency duration tracking responses in individual subjects. **(A-E)** Scalp topography of the high frequency duration-tracking score and single-electrode time-frequency plots for five individual subjects in experiment 2. The left column shows topographies and plots for lower left visual field stimuli, and the right column for lower right visual field stimuli. Within each column and panel, the left figure shows the topography of the high-frequency duration-tracking score, with thin and thick outlines indicating significance before and after Bonferroni correction for 19 posterior region-of-interest electrodes. The duration-tracking analysis was carried out for each individual subject and electrode using variance across trials. The right figure shows time-frequency plots for 300, 900, and 1500 ms stimulus duration, from top to bottom.

**Figure S4:**
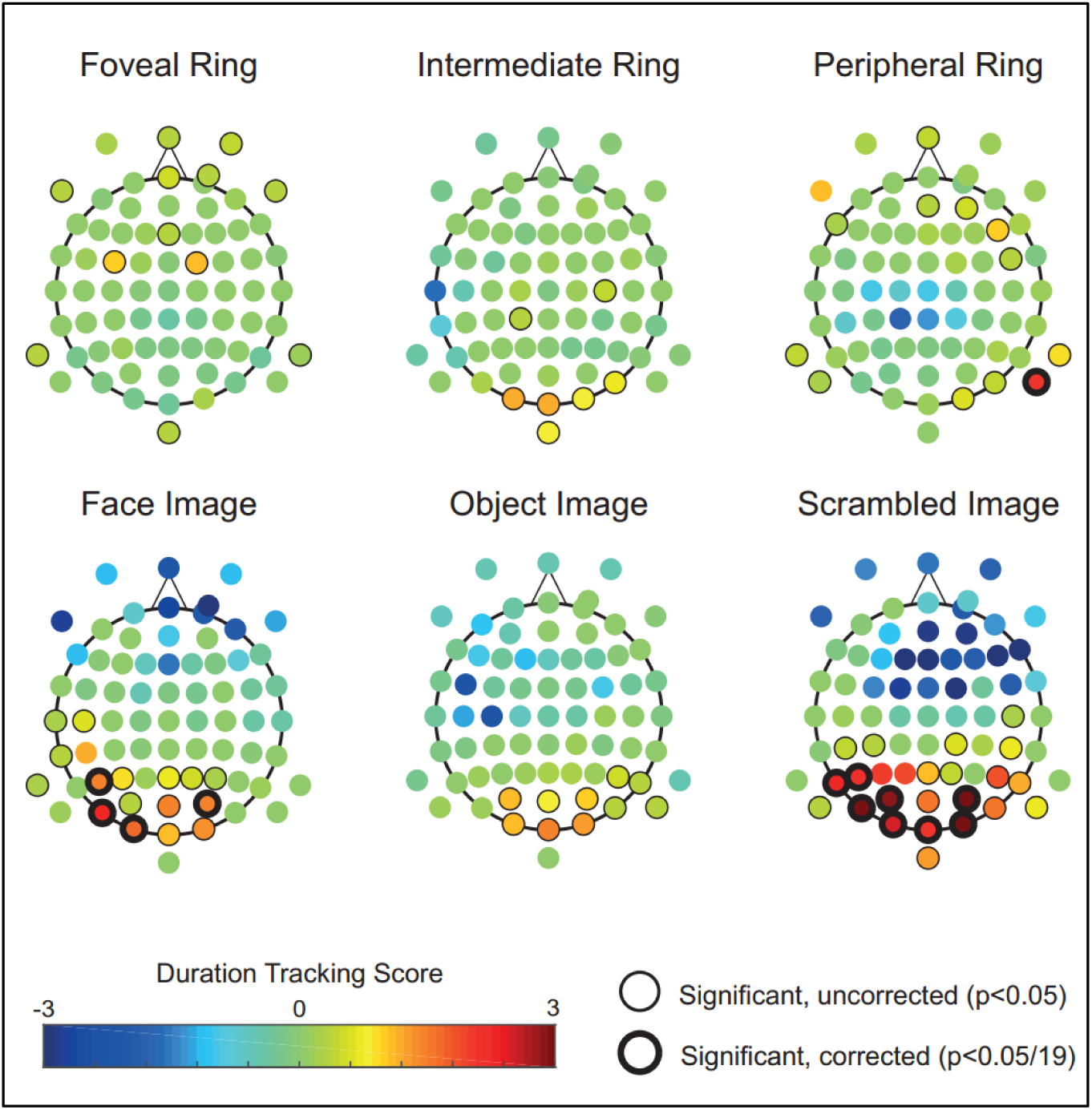
Experiment 3 – ERP duration-tracking by condition. Scalp topography of the group-average duration-tracking baseline shift effect, for each stimulus condition in experiment 3 (central, intermediate and peripheral checkerboard rings, and face, object and scrambled Mooney images).

**Figure S5:**
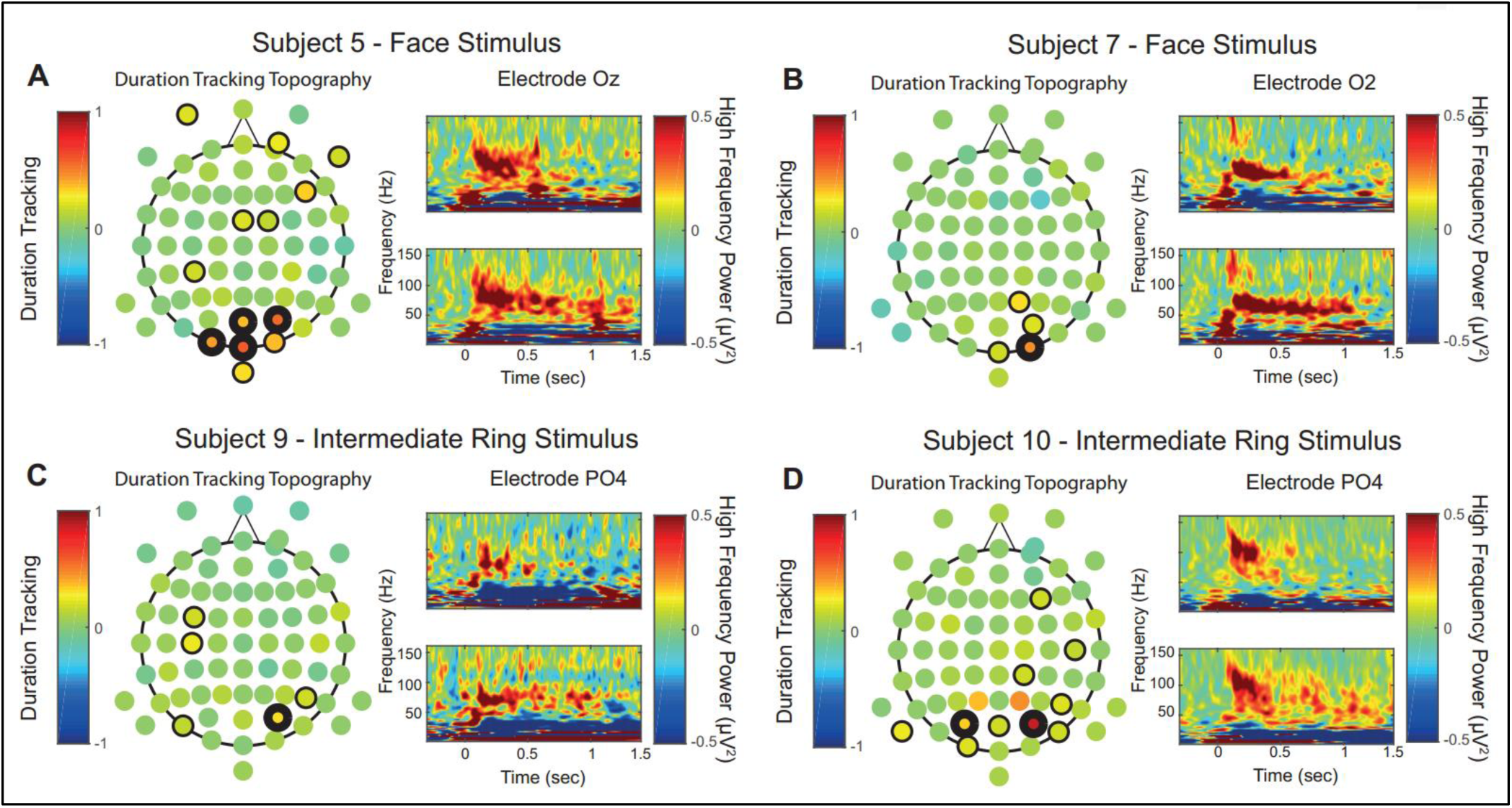
Experiment 3 – examples of high frequency duration-tracking responses in individual subjects. Four of fourteen subjects had significant high-frequency duration-tracking responses in at least one condition. Subjects 5,7 and 9 had significant responses in three conditions, though only one condition per subject is shown in the figure. **(A-D)** Left: topography of the duration-tracking effect in one example condition. Thin outlines indicate p<0.05, thick outlines indicate p<0.05/19 (Bonferroni-corrected). Right: Time frequency plots for the 0.5 sec (top) and 1 sec (bottom) duration trials from the same condition, in a single electrode with a significant (corrected) effect.

